# Butyrate extends health and lifespan in mice with mitochondrial deficiency

**DOI:** 10.64898/2026.01.13.699287

**Authors:** Enrique Gabandé-Rodríguez, Manuel M. Gómez de las Heras, Pablo Ramírez-Ruiz de Erenchun, Carolina Simó, Virginia García-Cañas, Naohiro Inohara, Inés Berenguer-López, Violeta Enríquez-Zarralanga, Álvaro Fernández-Almeida, Jorge Oller, Gonzalo Soto-Heredero, Elisa Carrasco, Sandra Delgado-Pulido, José Ignacio Escrig-Larena, Isaac Francos-Quijorna, Raquel Justo-Méndez, Juan Francisco Aranda, Joanna Poulton, Ana Victoria Lechuga-Vieco, José Antonio Enríquez, Gabriel Núñez, María Mittelbrunn

**Author notes:** These authors contributed equally to this work (Enrique Gabandé-Rodríguez, Manuel M. Gómez de las Heras).

## Abstract

Mitochondrial diseases progressively lead to multisystemic failure with treatment options remaining extremely limited. To investigate novel strategies that alleviate mitochondrial dysfunction, we have generated an ubiquitous and tamoxifen-inducible knockout mouse model of mitochondrial transcription factor A (TFAM), a nuclear-encoded protein involved in mitochondrial DNA (mtDNA) maintenance — *Tfam*^fl/fl^*Ub*^Cre-ERT2^ (iTfamKO) mice. Systemic TFAM deficiency triggers mitochondrial decline in a myriad of tissues in adult mice. Consequently, iTfamKO mice manifest multiorgan dysfunction including lipodystrophy, sarcopenia, metabolic alterations, kidney failure, neurodegeneration, and locomotor dysregulation, which result in the premature death of these mice. Interestingly, iTfamKO mice display intestinal barrier disruption and gut dysbiosis, with diminished levels of microbiota-derived short-fatty acids (SCFAs), such as butyrate. Mice with a deficient proof-reading version of the mtDNA polymerase gamma (mtDNA-mutator mice) phenocopy the dysfunction of the intestinal barrier and bacterial dysbiosis with reduced levels of butyrate, suggesting that different mouse models of mitochondrial dysfunction share deficient generation of butyrate. Transfer of microbiota from healthy control mice or administration of tributyrin, a butyrate precursor, delay multiple signs of multimorbidity extending lifespan in iTfamKO mice. Mechanistically, butyrate supplementation recovers epigenetic histone acylation marks that are lost in the intestine of *Tfam* deficient mice. Overall, our findings highlight the relevance of preserving host-microbiota symbiosis in disorders related to mitochondrial dysfunction.

## Introduction

Mitochondria are highly dynamic organelles that serve as bioenergetic and signaling hubs in eukaryotic cells supporting relevant functions including calcium handling, regulation of apoptosis and oxidative stress, lipid biosynthesis and, more importantly, production of the main cellular fuel — ATP. One of the peculiarities of mitochondria is that these organelles harbor their own genome, the mitochondrial DNA (mtDNA), which encodes 13 subunits of the electron transport chain that supports oxidative phosphorylation (OXPHOS) for ATP production^1^. Replication, transcription, and maintenance of mtDNA are controlled by the nuclear-encoded mitochondrial transcription factor A (TFAM)^2^. TFAM is essential for OXPHOS, mitochondrial biogenesis, and embryonic development, and the generation of tissue-specific knockout mouse models have further illustrated the importance of mitochondria in different cell types^3-12^. Consistently, defects impinging on mitochondrial function precipitate the development of mitochondrial diseases, a group of inherited metabolic disorders showing severe muscular, cardiovascular, metabolic, and neurologic manifestations^13,14^. Although guidelines are available in the clinic to manage symptoms and complications of mitochondrial disease, limited therapeutic options are yet accessible^15^.

The intestine is a biological barrier regulated by an intimately attached epithelial cell layer that secretes mucus and antimicrobial peptides. Importantly, the collapse of this barrier is associated with increased morbidity and mortality in different species^16-18^. Beyond energy supply, mitochondrial metabolism plays a major role in gut homeostasis since it coordinates intestinal stem cell self-renewal and differentiation, cell death programs, smooth muscle contractility, and oxygen bioavailability in the lumen^19^. Accordingly, patients living with mitochondrial disorders usually suffer from chronic mucosal atrophy and gastrointestinal dysmotility with delayed emptying resulting in severe constipation^20^. In addition, the intestine harbors a highly diverse ecosystem of microorganisms, the gut microbiota, which lives in symbiosis with the host providing pivotal metabolic pathways. Among others, short chain fatty acids (SCFAs) synthesized by intestinal bacteria such as acetate, propionate, and butyrate strengthen the gut barrier and shape host physiology^21^. Mechanistically, SCFAs can trigger extracellular and intracellular signaling cascades as well as remodel the epigenetic landscape, therefore fine-tuning immunity, metabolism, and gene expression in the host^22-25^. Lately, research has shown that microbiota-derived butyrate regulates gene expression in the intestine of mice by controlling histone H3 butyrylation levels^26^. Given their pleiotropic effects, manipulation of SCFAs has been proposed as therapeutic option for a myriad of inflammatory, cardiometabolic, and neurologic diseases^27^. While these results seem encouraging, the effectiveness of SCFAs in models of mitochondrial disease that display systemic multiorgan failure remains uncertain. Herein, we have explored this line of intervention in a novel mouse model of mitochondrial dysfunction by inducible, and ubiquitous deletion of TFAM, which mirrors the multimorbidity associated with mitochondrial diseases.

## Results

### Deletion of TFAM in adult mice induces systemic organ failure

To induce mitochondrial decline in adult mice, we crossed mice carrying loxP-flanked alleles of the gene encoding TFAM (*Tfam*^fl/fl^) with mice expressing a tamoxifen-inducible Cre-ERT2 recombinase under the control of the ubiquitous *Ubc* gene (*Ubc*^Cre-ERT2^) to generate *Tfam*^fl/fl^*Ubc*^Cre-ERT2^ mice. This system enabled systemic, and inducible knockdown of *Tfam* gene in adult mice (**Figure S1A**), avoiding the reported embryonic lethality of germline TFAM deletion^2^. Consistently, we observed a remarkable drop in the levels of *Tfam* transcripts in a myriad of tissues following tamoxifen administration (**Figure S1B**). Over time, adult *Tfam*^fl/fl^*Ubc*^Cre/ERT2+/–^ mice (henceforth iTfamKO mice) manifested a progeroid phenotype starting 70 to 90 days after tamoxifen administration, with a shortened lifespan compared with age-matched control littermates (**Figures 1A and 1B**).

**Figure 1.**
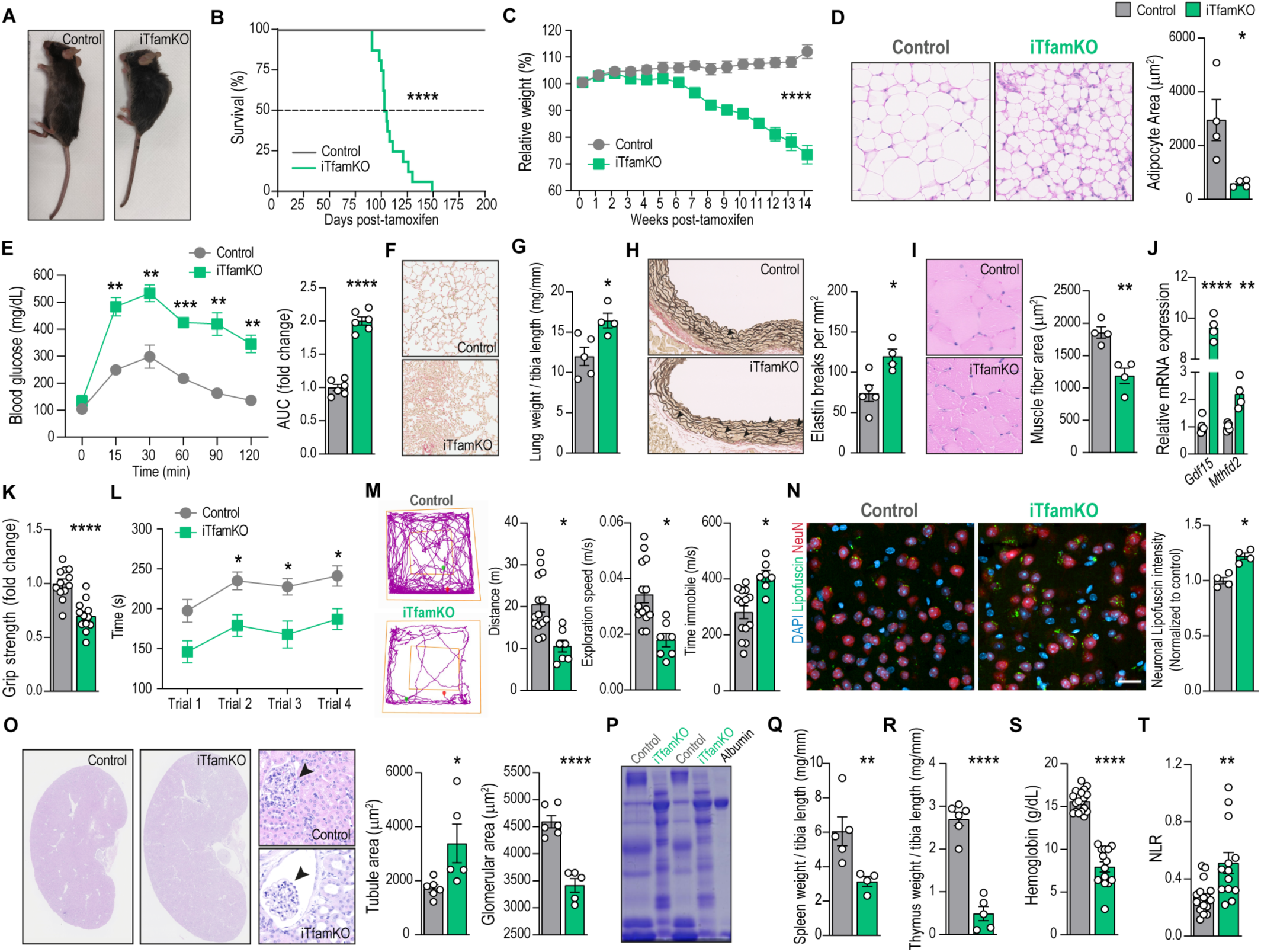
Deletion of *Tfam* in adult mice induces systemic organ failure. (**A**) Representative picture of control and iTfamKO mice 90 days after tamoxifen administration. (**B**) Longitudinal assessment of relative body weight following tamoxifen administration (*n* = 15 to 17). (**C**) Kaplan-Meier survival curves (*n* = 7 to 16). (**D**) Representative hematoxylin and eosin (H&E)–stained sections of gonadal white adipose tissue and quantification of adipocyte surface area (*n* = 4). (**E**) Glucose tolerance tests and its respective area under the curve (AUC) (*n* = 6). (**F**) Representative Sirius Red–stained sections of lung parenchyma. (**G**) Quantification of lung weight normalized to tibia length (*n* = 4 to 5). (**H**) Representative elastic van Gieson (EVG)-stained sections of aorta and quantification of elastin breaks (*n* = 4 to 5). (**I**) Representative H&E–stained sections of skeletal muscle and quantification of myofiber cross-sectional area (*n* = 4). (**J**) Relative mRNA levels of the mitochondrial disease-associated genes *Gdf15* and *Mthfd2* in the skeletal muscle (*n* = 4 to 5 mice). (**K**) Forelimb grip strength test (*n* = 12). (**L**) Rotarod test performance (*n* = 20). (**M**) Open field test performance (*n* = 7 to 14). (**N**) Representative image (scale bar, 20 μm) and quantification of lipofuscin particles in brain neurons (*n* = 4). (**O**) Representative H&E–stained sections of kidney parenchyma and quantification of glomerular and tubule area (*n* = 5 to 6). (**P**) Representative Coomassie-stained gel of urine samples. (**Q** and **R**) Quantification of (Q) spleen and (R) thymus weight normalized to tibia length (*n* = 4 to 6). (**S**) Concentration of hemoglobin (*n* = 14 to 18). (**T**) Neutrophil-to-lymphocyte ratio (NLR) in the blood (*n* = 13 to 17). Data are shown as means ± SEM, where each dot is a biological sample. *P* values were determined by (B) log-rank (Mantel-Cox) test, (C) two-way analysis of variance (ANOVA), (D, G to K, M to O, and Q to S) unpaired Student’s *t* test, (L) mixed-effects analysis with Šidák’s multiple comparisons test, or (T) two-tailed Mann–Whitney *U* test. (E) *P* values were determined by (curve) two-way ANOVA with Šidák’s multiple comparisons test or (AUC) unpaired Student’s *t* test. \**P* ≤ 0.05; \*\**P* ≤ 0.01; \*\*\**P* ≤ 0.001; and \*\*\*\**P* ≤ 0.0001.

In addition, these mice developed a multisystemic morbidity syndrome that encompasses a marked loss of body weight and lipodystrophy, showing an acute reduction of white and brown adipose tissues as well as smaller adipocytes (**Figures 1C and 1D and Figures S1C and S1D**). This was accompanied by diminished body temperature (**Figure S1E**) and signs of glucose intolerance (**Figure 1E**). iTfamKO mice also manifested hypogonadism, namely a reduced size of both testicles in males and ovaries in females (**Figure S1F**). Regarding the cardiopulmonary system, iTfamKO mice displayed severe lung fibrosis (**Figure 1F**) with increased lung weight, suggesting lung congestion or pulmonary oedema (**Figure 1G**). Histological evaluation of aortas revealed an increase in aortic dissections (**Figure 1H**), in line with previous works of smooth muscle cell-specific deletion of *Tfam*^7^. Furthermore, analysis of skeletal muscle in iTfamKO mice uncovered smaller diameter of muscle fibers and increased expression of growth differentiation factor 15 (*Gdf15*) and methylene tetrahydrofolate dehydrogenase 2 (*Mthfd2*), biomarkers of mitochondrial disease^28^ which, together with a lower grip strength, were suggestive of severe sarcopenia (**Figures 1I-K**). iTfamKO mice also exhibited signs of neurological disability, such as impaired locomotor coordination and activity in the rotarod and Open field tests (**Figures 1L and 1M and Figure S1G**), as well as reduced performance in the nest building test (**Figures S1H**). In addition, the brain of iTfamKO mice was smaller (**Figure S1I**) and harbored markers of neurodegeneration, such as increased neuronal lipofuscin aggregates (**Figure 1N**), reduced levels of the postsynaptic marker PSD-95, and increased levels of phosphorylated Tau, as a consequence of successful depletion of *Tfam* in the brain (**Figures S1J and S1K**).

Importantly, iTfamKO mice displayed kidney failure with abnormally enlarged kidneys featuring signs of glomerulosclerosis and kidney tubule dilation (**Figure 1O**), which was associated with polyuria and extremely clear urine, commonly found in diabetic conditions, as well as albuminuria (**Figures 1P and Figure S1L**). Furthermore, we observed that iTfamKO mice displayed premature inflammaging including heightened levels of interleukin (IL)-6 and tumor necrosis factor (TNF) in the serum (**Figure S1M**). Similarly, the liver of iTfamKO mice showed enhanced expression of proinflammatory markers like IL-6 and TNF and downregulation of anti-inflammatory molecules like IL-10, together with strong expression of pro-fibrotic genes such as *Serpin-1*, *Ccn2*, and *Spp1* (**Figure S1N**). Of note, this was accompanied by increased expression of the senescence markers *Cdkn2a* and *Cdkn1a*, respectively encoding the cyclin-dependent kinase inhibitors P16^INK4a^ and P21^Waf/Cip1^, in the liver of iTfamKO mice (**Figure S1N**).

Lastly, the spleen and thymus of iTfamKO mice showed a substantial atrophy (**Figures 1Q and 1R**), indicative of alterations in the hematopoietic system. Accordingly, hematological analysis of iTfamKO mice revealed decreased levels of parameters related to the number and function of erythrocytes (**Figure S2A**), highlighting a profile of severe anemia (**Figures 1S**). We also observed thrombocytosis and variations in leukocytes including lymphopenia and neutrophilia leading to a heightened neutrophil-to-lymphocyte ratio (NLR), a parameter predicting poor survival in multiple diseases^29^, in iTfamKO mice compared with controls (**Figure 1T** and **Figures S2B and S2C**).

These data demonstrate that systemic ablation of *Tfam* in adult mice drives systemic organ failure including sarcopenia, cardiometabolic alterations, anemia, renal failure, and neurodegeneration, concomitant with the induction of a pro-inflammatory and senescence program in several tissues.

### Intestinal barrier integrity is disrupted in iTfamKO mice

Given the importance of mitochondrial fitness in maintaining intestinal barrier integrity^19^, whose breakdown is associated with a plethora of pathological conditions^30^, we sought to investigate the status of the small and large intestine of adult iTfamKO mice. We confirmed that the expression of *Tfam* in both the small intestine and colon was notably diminished 90 days upon the induction of the Cre recombinase expression (**Figure 2A**). Accordingly, electron microscopy analysis uncovered that mitochondria in the small intestine and colon of iTfamKO mice were smaller and showed a remarkable increase in electrodense inclusion bodies (**Figures 2B and 2C**), which were compatible with previously reported abnormal calcium phosphate aggregates due to a faulty handling of calcium^31^. Macroscopic examination of these organs revealed a significant increase in the length of the small intestine, but not in the colon, in iTfamKO mice compared with controls (**Figure 2D**). Moreover, histological evaluation of the small intestine showed an altered tissue architecture with notably smaller crypts (**Figures 2E and 2F**). These alterations were concomitant with a decreased number of proliferative Ki67^+^ cells in the crypts (**Figures 2G and 2H**), and enhanced expression of the senescence marker P21^Waf/Cip1^ in the small intestine of iTfamKO mice compared with control littermates (**Figure 2I**).

**Figure 2.**
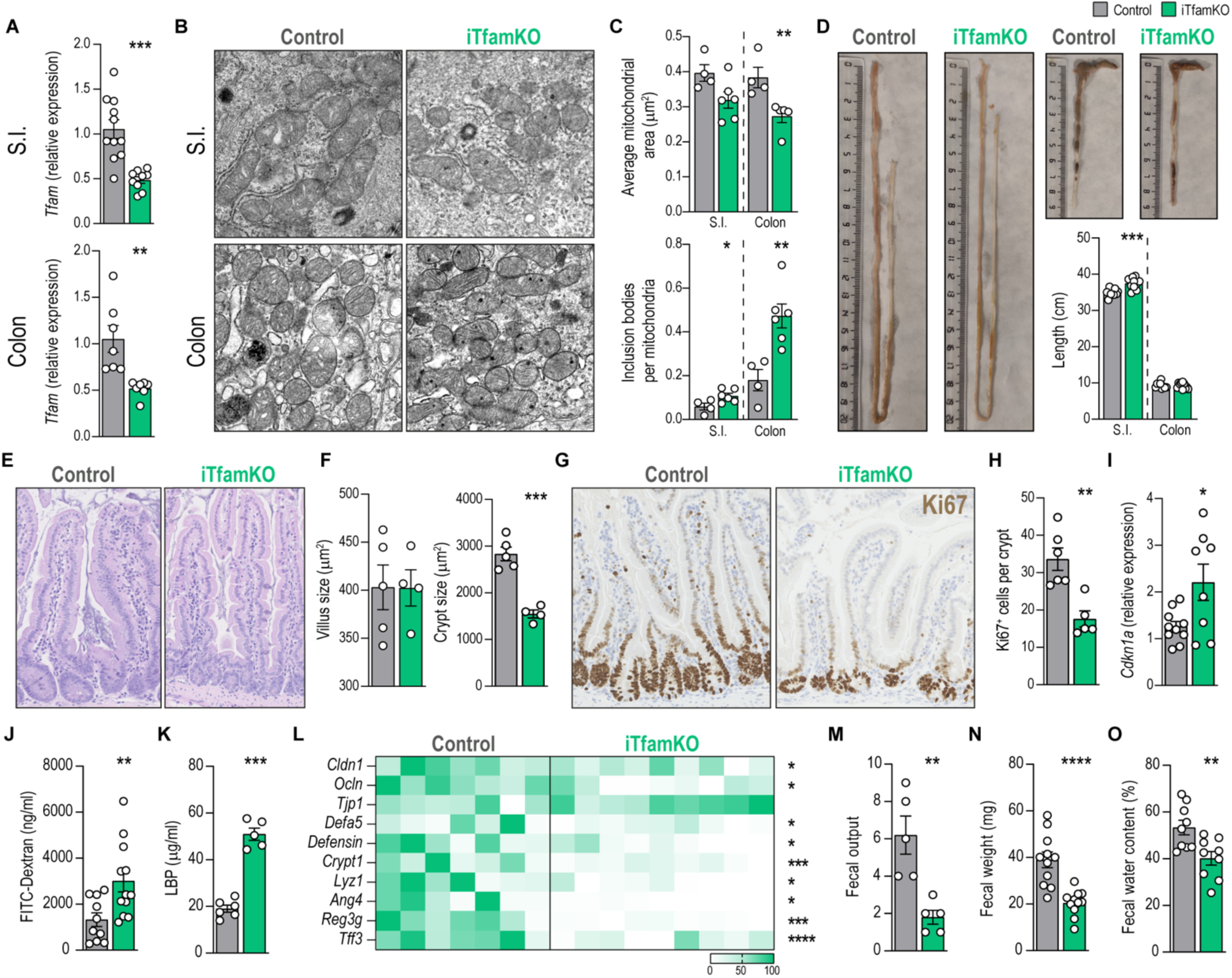
iTfamKO mice display intestinal barrier disruption and severe constipation. (**A**) Relative expression levels of *Tfam* in the small intestine (S.I.) and the colon of control and iTfamKO mice 90 days after tamoxifen administration (*n* = 7 to 11). (**B**) Representative electron microscopy images of the S.I. and the colon epithelium. (**C**) Quantification of mitochondrial area and the number of electrodense inclusion bodies per mitochondria in the S.I. and the colon epithelium (*n* = 4 to 6). (**D**) Representative picture of the S.I. and the colon, and its respective length quantification (*n* = 9 to 10). (**E**) Representative hematoxylin and eosin (H&E)–stained sections of the S.I. (**F**) Quantification of villus size and crypt area in the S.I. (*n* = 4 to 5). (**G**) Representative immunohistochemistry staining of the cell proliferation marker Ki67 in the S.I. (**H**) Quantification of Ki67-positive cells per crypt in the S.I. (*n* = 5 to 6). (**I**) Relative mRNA levels of the senescence-associated gene *Cdkn1a* in the S.I. (*n* = 9 to 10). (**J**) Concentration of FITC-dextran in the serum (*n* = 10 to 12). (**K**) Levels of LPS-binding protein (LBP) in the serum (*n* = 5 to 6). (**L**) Heatmap depicting normalized values of qPCR analysis of genes associated with tight junctions (*Cldn1*, *Ocln*, *Tjp1*), antimicrobial peptides (*Defa5*, *Defensin*, *Crypt1*, *Lyz1*, *Ang4*, *Reg3g*), and barrier function (*Tff3*) in the S.I. (*n* = 7 to 9 mice). (**M**) Quantification of the number of feces defecated in 30 min (*n* = 5). (**N**) Quantification of feces weight (*n* = 11). (**O**) Percentage of water in the feces (*n* = 9). Data are shown as means ± SEM, where each dot is a biological sample. *P* values were determined by unpaired Student’s *t* test. \**P* ≤ 0.05; \*\**P* ≤ 0.01; \*\*\**P* ≤ 0.001; and \*\*\*\**P* ≤ 0.0001.

Furthermore, intestinal barrier integrity is severely compromised in this mouse model, as confirmed by increased penetration of FITC-dextran after oral administration (**Figure 2J**). To test whether gut hyperpermeability resulted in a heightened translocation of bacteria to the periphery, we quantified the amount of LPS-binding protein (LBP) in serum of these mice, a surrogate marker of bacterial translocation that dramatically increased in iTfamKO mice (**Figure 2K**). We further assessed the physical integrity of the intestinal barrier by evaluating markers of cell-to-cell junctional complexes in the intestine, for instance expression of genes encoding proteins involved in tight junctions (*Cldn1*, *Ocln*), antimicrobial peptides (*Defa5*, *Defensin*, *Crypt1*, *Lyz1*, *Ang4*, and *Reg3g*) and participating in mucosa regeneration (*Tff3*), which showed an overall downregulation in the small intestine, but not in the colon, of iTfamKO mice compared with controls (**Figures 2L and Figure S3**). We did not observe any traits of colitis such as shortened colon or blood in the feces of iTfamKO mice. Instead, these mice exhibited marked signs of constipation, as demonstrated by a decrease in the frequency of defecated stool samples (**Figure 2M**), together with a substantially reduced weight and percentage of water content in feces (**Figures 2N and 2O**), suggesting defects in gut peristalsis and water absorption.

### Systemic deficiency of *Tfam* leads to bacterial dysbiosis and reduced SCFA levels

Metabolic perturbations in the intestine are associated with pathogenic shifts in the gut microbiota^32,33^. To characterize the composition of the bacterial communities in the intestine of iTfamKO mice, we performed *16S* rRNA gene sequencing in samples of the terminal ileum and the colon in these mice. Analyses of α-diversity indicated a reduction in parameters such as Shannon and observed species (Sobs) indexes in the ileal but not in the colonic microbiota (**Figures 3A and 3B**). Moreover, β-diversity analyses by non-metric multidimensional scaling (NMDS) denoted a remarkably distinct configuration of the gut microbiota in the colon of iTfamKO mice compared with control littermates (**Figures 3C and 3D**). Compositional analysis in both intestinal compartments suggested a striking reduction in bacteria belonging to Clostridiales order such as *Lachnospiraceae* and *Ruminococcaceae* families in the ileal and colonic microbiome of iTfamKO mice when compared with controls (**Figure 3E**). These changes were accompanied by an expansion of bacteria from the Bacillales order, including members of the *Staphylococcaceae* family in both the ileum and colon of this mouse model (**Figure 3E**).

**Figure 3.**
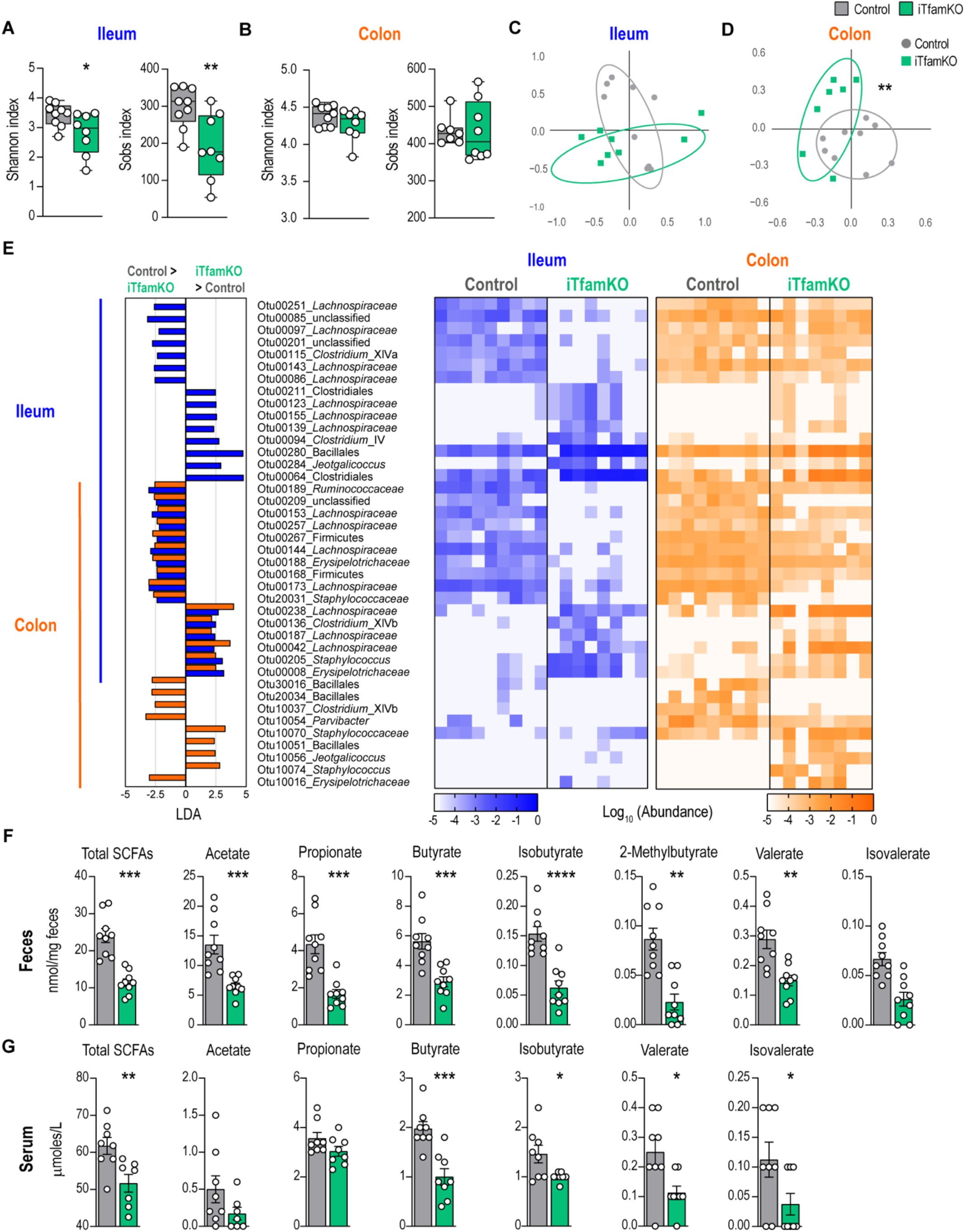
Systemic deficiency of *Tfam* leads to bacterial dysbiosis and reduced SCFA production. (**A** and **B**) Shannon index and observed species index (Sobs) parameters of α-diversity in (A) the ileum and (B) colon-resident microbiota of control and iTfamKO mice 90 days after tamoxifen administration (*n* = 8 to 9). (**C** and **D**) Non-metric multidimensional scaling (NMDS) plots of β-diversity values (θYC indexes) in (C) the ileum and (D) colon-resident microbiota (*n* = 8 to 9). (**E**) Left: differentially abundant operational taxonomic units (OTUs) depicted with linear discriminant analysis (LDA) values of linear discriminant effect size (LEfSe, *P* < 0.05; FDR, *Q* < 0.05; |SNR| > 0.5; |LDA| > 2; |fold change| > 10; maximal abundance > 0.001) comparing ileal and colonic microbiota in control versus iTfamKO mice. Right: heatmap depicting abundance values. (**F** and **G**) Quantification of short-chain fatty acids (SCFAs) in (F) the feces and (G) the serum (*n* = 8 to 9). Data are shown as means ± SEM, where each dot is a biological sample. *P* values were determined by (A, C, F, and G) unpaired Student’s *t* test or (B and D) permutational multivariate analysis of variance (PERMANOVA). \**P* ≤ 0.05; \*\**P* ≤ 0.01; \*\*\**P* ≤ 0.001; and \*\*\*\**P* ≤ 0.0001.

To delve into the *16S* rRNA metagenomic data, we performed a predictive functional analysis of the gut microbiome. Notably, the sequencing profiles of the ileum-dwelling microbiota unveiled a reduction in biochemical pathways related to carbohydrate degradation, for instance, fucose, D-glucarate, and rhamnose in iTfamKO mice (**Figure S4A**). Polysaccharide utilization by the gut microbiota leads to the balanced production of health-promoting metabolites such as SCFAs ^34^. To examine whether the observed compositional and metabolic changes of the gut microbiota impinged on the production of SCFAs, we quantified these molecular species in fecal and serum samples of mice using liquid chromatography-mass spectrometry (LC-MS). Remarkably, we observed a reduction in the concentration of total SCFAs, as well as in each one of them (i.e., acetate, propionate, butyrate, isobutyrate, 2-methylbutyrate, valerate, and isovalerate) in the feces as well as in the serum of iTfamKO mice when compared with control littermates (**Figure 3F and 3G**), supporting that these alterations also occur systemically.

In an effort to shed light on which bacteria could be involved in this drop of SCFAs, we performed Spearman correlation analyses between the ileum and colon-resident microbiota and the concentration of SCFA species in the same samples. Focusing on the operational taxonomic units (OTUs) that decreased in the gut microbiota of iTfamKO mice, the results unveiled ten OTUs that positively correlated with the reduced levels of these molecules (**Figure S4B**). We found among them members belonging to the *Ruminococcaceae* and *Lachnospiraceae* families, which represent crucial SCFA-producing bacteria in the gut microbiota.

Altogether, our findings suggest that deletion of *Tfam* leads to mitochondrial alterations in the intestine, resulting in the rupture of the intestinal barrier, gut dysmotility, and bacterial dysbiosis that is associated with a marked reduction in SCFA production.

### mtDNA-mutator mice display gut dysbiosis and reduced levels of butyrate

We wondered whether another mouse model of mitochondrial deficiency showing premature multimorbidity also exhibited alterations in intestinal barrier integrity and gut microbiota composition. For that, we examined *PolgA*^D257A/D257A^ (*Polg*^mut^) or mtDNA-mutator mice that carry a deficient proof-reading version of the gene encoding the mtDNA polymerase gamma, therefore leading to a progressive accumulation of somatic mutations in the mtDNA that trigger premature multimorbidity^35,36^. We analyzed *Polg*^mut^ mice at 50 weeks of age, once they show signs of multimorbidity, compared with age-matched control mice (**Figure 4A**).

**Figure 4.**
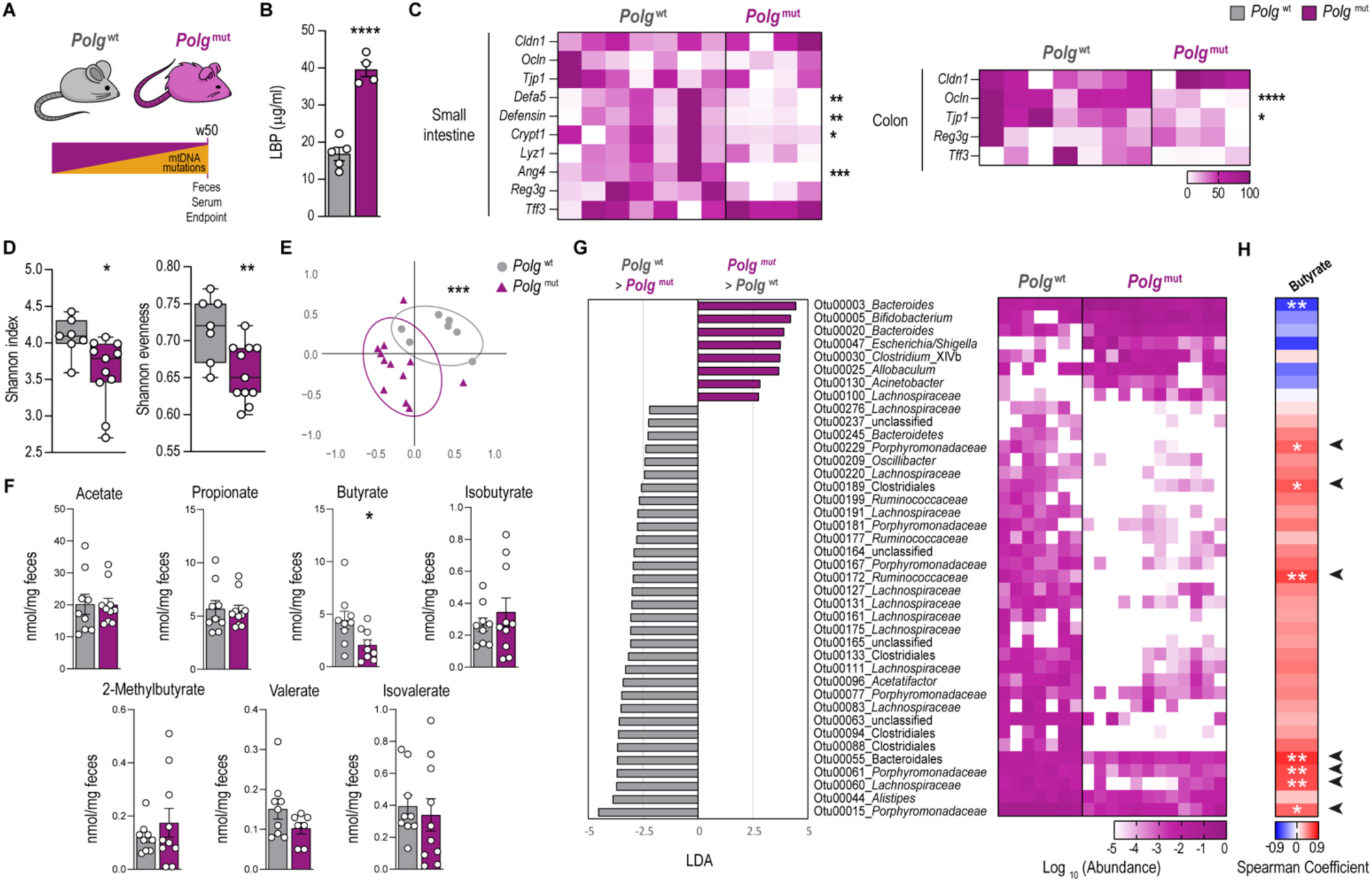
mtDNA-mutator mice show bacterial dysbiosis and reduced levels of butyrate. (**A**) Scheme of 50-week-old *Polg*^wt^ and *Polg*^mut^ mice. (**B**) Levels of LPS-binding protein (LBP) in the serum (*n* = 4 to 5). (**C**) Heatmap depicting normalized values of qPCR analysis of genes associated with tight junctions (*Cldn1*, *Ocln*, *Tjp1*), antimicrobial peptides (*Defa5*, *Defensin*, *Crypt1*, *Lyz1*, *Ang4*, *Reg3g*), and barrier function (*Tff3*) in the small intestine and the colon (*n* = 4 to 7). (**D**) Shannon index and Shannon evenness parameters of α-diversity in fecal microbiota (*n* = 7 to 11). (**E**) Non-metric multidimensional scaling (NMDS) plots of β-diversity values (θYC indexes) in fecal microbiota (*n* = 7 to 12). (**F**) Quantification of short-chain fatty acid species in the feces (*n* = 9 to 11). (**G**) Left: differentially abundant operational taxonomic units (OTUs) depicted with linear discriminant analysis (LDA) values of linear discriminant effect size (LEfSe, *P* < 0.05; FDR, *Q* < 0.05; |SNR| > 0.5; |LDA| > 1; |fold change| > 5; maximal abundance > 0.001) comparing fecal microbiota in *Polg*^wt^ versus *Polg*^mut^ mice. Right: heatmap depicting abundance values. (**H**) Heatmap depicting Spearman’s rank correlation coefficients between differentially abundant OTUs and the concentration of butyrate in the feces. Data are shown as means ± SEM, where each dot is a biological sample. *P* values were determined by (B to D, and F) unpaired Student’s *t* test or (E) permutational multivariate analysis of variance (PERMANOVA). \**P* ≤ 0.05; \*\**P* ≤ 0.01; \*\*\**P* ≤ 0.001; and \*\*\*\**P* ≤ 0.0001.

Notably, *Polg*^mut^ mice displayed intestinal barrier dysfunction with 2-fold heightened levels of LBP in the serum compared with control littermates (**Figure 4B**). This was accompanied by downregulated expression of genes encoding antimicrobial peptides such as *Defa5*, *Defensin*, *Crypt*, and *Ang4* in the small intestine, and reduced expression of genes participating in tight junction architecture like *Ocln* and *Tjp1* in the colon of *Polg*^mut^ mice compared with controls (**Figure 4C**). Analyses of the fecal microbiota in these mice unveiled a marked reduction in α-diversity parameters such as Shannon index and Shannon evenness in *Polg*^mut^ mice compared with controls (**Figure 4D**), suggesting a loss of bacterial species in this mouse model. In addition, analyses of β-diversity by NMDS demonstrated a substantially different configuration of the fecal microbiota (**Figure 4E**). Similar to what observed in iTfamKO mice, LC-MS quantification of SCFA species in the feces of *Polg*^mut^ mice uncovered decreased levels of butyrate in these mice when compared to control mice (**Figure 4F**). Notably, predictive functional analysis of the *16S* rRNA metagenomic data showed downregulation of biochemical pathways related to carbohydrate utilization (**Figure S5**). In addition, evaluation of predictive metabolic pathways shared by iTfamKO and *Polg*^mut^ mice showed downregulation in pathways linked to the degradation of aromatic compounds (**Figure S6**), which are usually metabolized into acetyl-Coenzyme A (acetyl-CoA) — a substrate for butyrate biosynthesis in intestinal bacteria.

Evaluation of the relative abundance of bacteria in the feces of these mice showed an overall reduction in bacteria belonging to the Clostridiales order such as the *Lachnospiraceae* and *Ruminococcaceae* families and belonging to Bacteroidales including the *Porphyromonadaceae* family. We found an abnormal expansion of *Bacteroidaceae* and some members of *Enterobacteriaceae* in *Polg*^mut^ mice compared with control littermates (**Figure 4G**). Spearman correlation analysis between differentially abundant OTUs and butyrate levels revealed seven OTUs that were diminished in these mutant mice and positively correlated with the decreased levels of butyrate. These OTUs corresponded to well-known butyrate-producing members of the *Lachnospiraceae*, *Ruminococcaceae*, or *Porphyromonadaceae* families (**Figure 4H**).

Thereby, our findings reveal that an alternative mouse model of multimorbidity due to mitochondrial dysfunction manifests disruption of the gut barrier integrity as well as gut dysbiosis featured by the loss of butyrate-producing taxa.

### Butyrate supplementation improves systemic organ failure extending health and lifespan in iTfamKO mice

Given the intestinal dysbiosis found in iTfamKO mice, we investigated the contribution of the gut microbiota to the multimorbidity phenotype of these mice. Thus, we designed a strategy to replace the gut microbiota with fecal microbiota transplantation (FMT) assays. In brief, we transferred fecal microbiota from either control or iTfamKO donor mice to iTfamKO recipient mice (denoted as iTfamKO FMT_Control_ and iTfamKO FMT_KO_, respectively) starting 30 days after tamoxifen administration (**Figure 5A**). iTfamKO FMT_Control_ mice showed slightly delayed loss of body weight and increased muscle strength when compared with iTfamKO FMT_KO_ mice (**Figures 5B and 5C**), with no changes in the levels of LBP (**Figure 5D**). Moreover, water content in the feces was recovered in iTfamKO FMT_Control_ mice to levels of control mice (**Figure 5E**), suggesting improvement in signs of constipation. On top of this, we found that the levels of some SCFAs such as butyrate and, to a lesser extent, isobutyrate, 2-methylbutyrate, and isovalerate were restored in iTfamKO mice receiving microbiota from healthy control mice (**Figure 5F**). Notably, this was associated with an extension in the maximum lifespan by 70% (from 115 to 196 days) in iTfamKO FMT_Control_ mice compared with iTfamKO FMT_KO_ mice (**Figure 5G**).

**Figure 5.**
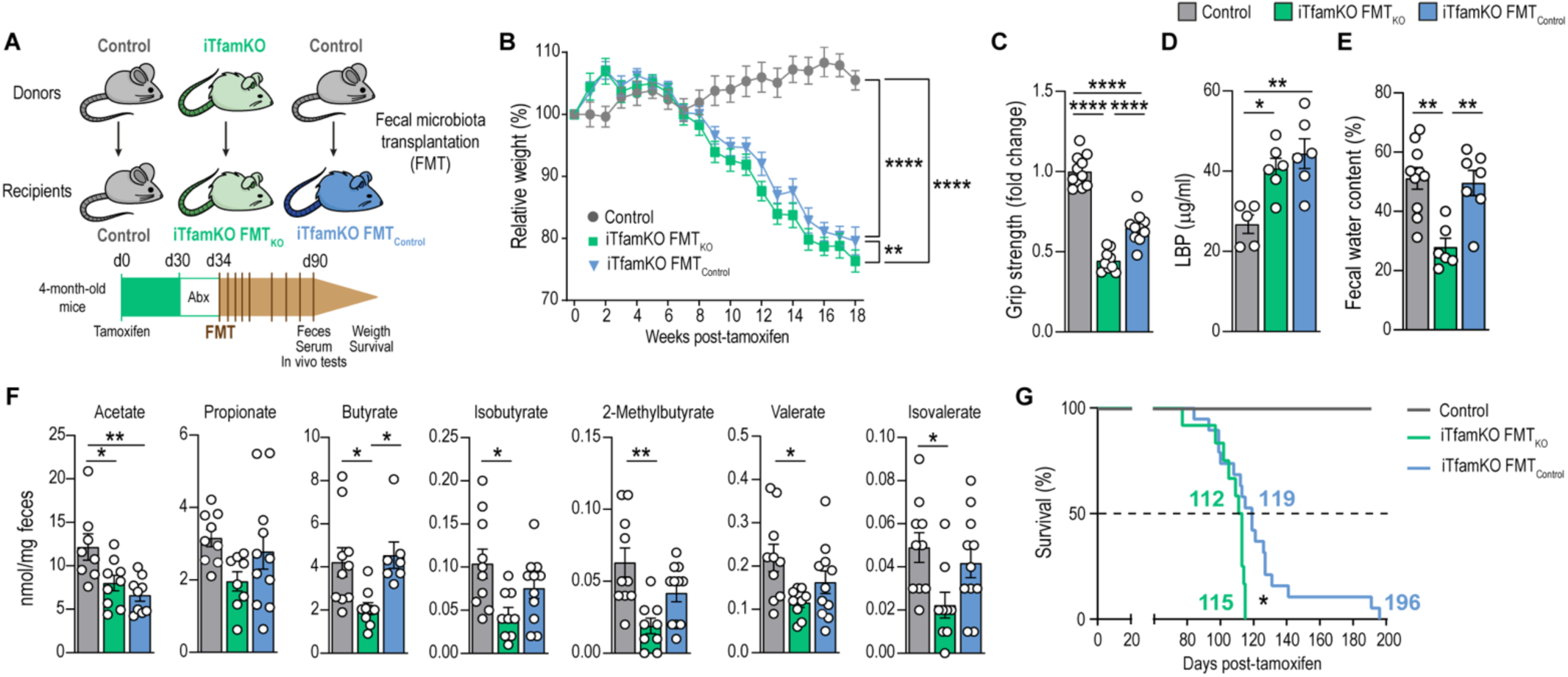
Transplantation of control microbiota ameliorates signs of multimorbidity in iTfamKO mice. (**A**) Experimental design of fecal microbiota transplantation (FMT) assays from donor control or iTfamKO mice into recipient iTfamKO mice (iTfamKO FMT_Control_ or iTfamKO FMT_KO_ mice, respectively). (**B**) Longitudinal assessment of body weight relative to tamoxifen administration (*n* = 19 to 23). (**C**) Forelimb grip strength analysis (*n* = 9 to 11). (**D**) Levels of LPS-binding protein (LBP) in the serum (*n* = 5 to 6). (**E**) Percentage of water in the feces (*n* = 6 to 10). (**F**) Quantification of short-chain fatty acid species in the feces (*n* = 8 to 11). (**G**) Kaplan-Meier survival curves (*n* = 9 to 19). Data are shown as means ± SEM, where each dot is a biological sample. *P* values were determined by (B) two-way analysis of variance (ANOVA) with Tukey’s multiple comparisons test, (C to F) one-way ANOVA with Tukey’s multiple comparisons test, or (G) log-rank (Mantel-Cox) test. \**P* ≤ 0.05; \*\**P* ≤ 0.01; and \*\*\*\**P* ≤ 0.0001.

Since FMT from healthy control mice rescued the levels of butyrate in iTfamKO recipient mice, being associated with improvement in healthspan and mortality, we further explored the therapeutic potential of restoring butyrate levels in this mouse model. For that purpose, iTfamKO mice were fed a diet supplemented with 10% tributyrin (TB), a butyrate precursor in the form of a triacyl-glycerol ester of butyric acids, starting 30 days after tamoxifen administration (**Figure 6A**). iTfamKO mice fed a TB-supplemented diet showed increased levels of butyrate in the feces compared with iTfamKO mice fed a standard diet (**Figures 6B and S7A**). Butyrate retrieval was associated with a delayed loss of body weight (**Figure 6C**), and notably enhanced muscle strength assessed by grip test, which was accompanied by diminished expression of the mitochondrial disease-associated genes *Gdf15* and *Mthfd2* in the skeletal muscle (**Figure 6D and 6E**). Furthermore, iTfamKO mice fed with a TB-supplemented diet showed improved glucose handling by glucose tolerance tests, and restored levels of fasting glucose almost to the levels of control mice (**Figure 6F and 6G**). Furthermore, quantification of albumin levels in the urine indicated improvement in the function of the kidney in iTfamKO mice fed a TB diet (**Figure 6H**). TB supplementation affected neither spleen size nor hematological parameters in treated iTfamKO mice (**Figures S7B and S7C**). Importantly, TB administration remarkably extended median lifespan by ∼ 25% (from 98 to 123 days) and maximum lifespan by more than 75% (from 106 to 186 days) in iTfamKO mice fed a TB diet when compared with those receiving standard diet (**Figure 6I**).

**Figure 6.**
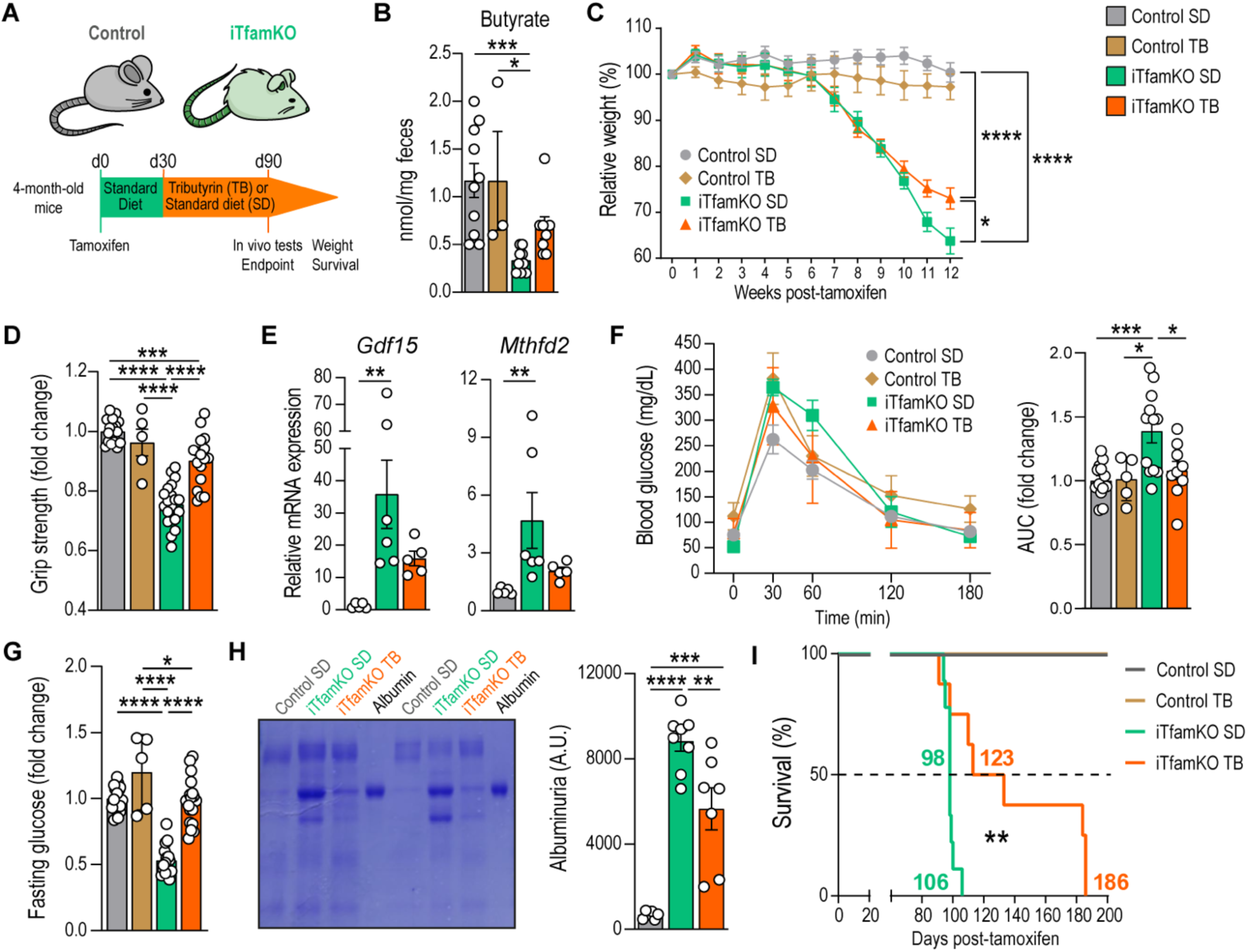
Tributyrin supplementation extends health and lifespan in iTfamKO mice. (**A**) Experimental design of control and iTfamKO mice fed either a standard diet (SD) or a 10% tributyrin-supplemented diet (TB). (**B**) Quantification of butyrate in the feces (*n* = 3 to 10). (**C**) Longitudinal assessment of body weight relative to tamoxifen administration (*n* = 5 to 21). (**D**) Forelimb grip strength analysis (*n* = 5 to 20). (**E**) Relative mRNA levels of the mitochondrial disease-associated genes *Gdf15* and *Mthfd2* in the skeletal muscle (*n* = 5 to 6 mice). (**F**) Glucose tolerance test and its respective area under the curve (AUC) quantification (*n* = 5 to 13). (**G**) Quantification of fasting glucose levels (*n* = 5 to 13). (**H**) Representative Coomassie-stained gel and quantification of urine samples (*n* = 5 to 9). (**I**) Kaplan-Meier survival curves (*n* = 5 to 10). Data are shown as means ± SEM, where each dot is a biological sample. *P* values were determined by (B and E) Kruskal-Wallis test with Dunn’s multiple comparisons test, (C) mixed-effects analysis with Šidák’s multiple comparisons test, (D, G, and H) one-way analysis of variance (ANOVA) with Tukey’s multiple comparisons test, or (I) log-rank (Mantel-Cox) test. (F) *P* values were determined by (curve) two-way ANOVA with Tukey’s multiple comparisons test or (AUC) one-way ANOVA with Tukey’s multiple comparisons test. \**P* ≤ 0.05; \*\**P* ≤ 0.01; \*\*\**P* ≤ 0.001; and \*\*\*\**P* ≤ 0.0001.

Taken together, our findings support the notion that interventions to restore butyrate levels delay signs of multimorbidity and extend the lifespan in iTfamKO mice.

### Butyrate retrieval restores epigenetic histone acylation marks in the intestine of iTfamKO mice

Gut microbiota-derived butyrate has been shown to remodel the epigenetic landscape of intestinal cells through histone H3 acylation^23,26^. To elucidate the molecular mechanisms underlying the beneficial effects of TB supplementation, we analyzed levels of acetylation and butyrylation on lysines 9 and 27 of histone H3 in the small intestine by Western Blotting. Compared with control mice, iTfamKO mice exhibited a notable reduction in the levels H3K9ac, H3K9bu, and H3K27bu in the small intestine (**Figure 7A**). Confirming the crucial role of microbiota-derived SCFAs in the acquisition of these histone modifications, we observed similar results in microbiota-depleted control mice treated with a cocktail of broad-spectrum antibiotics (Abx) (**Figure 7A**), which showed a near-complete loss of SCFAs compared with controls receiving vehicle (**Figure 7B**).

**Figure 7.**
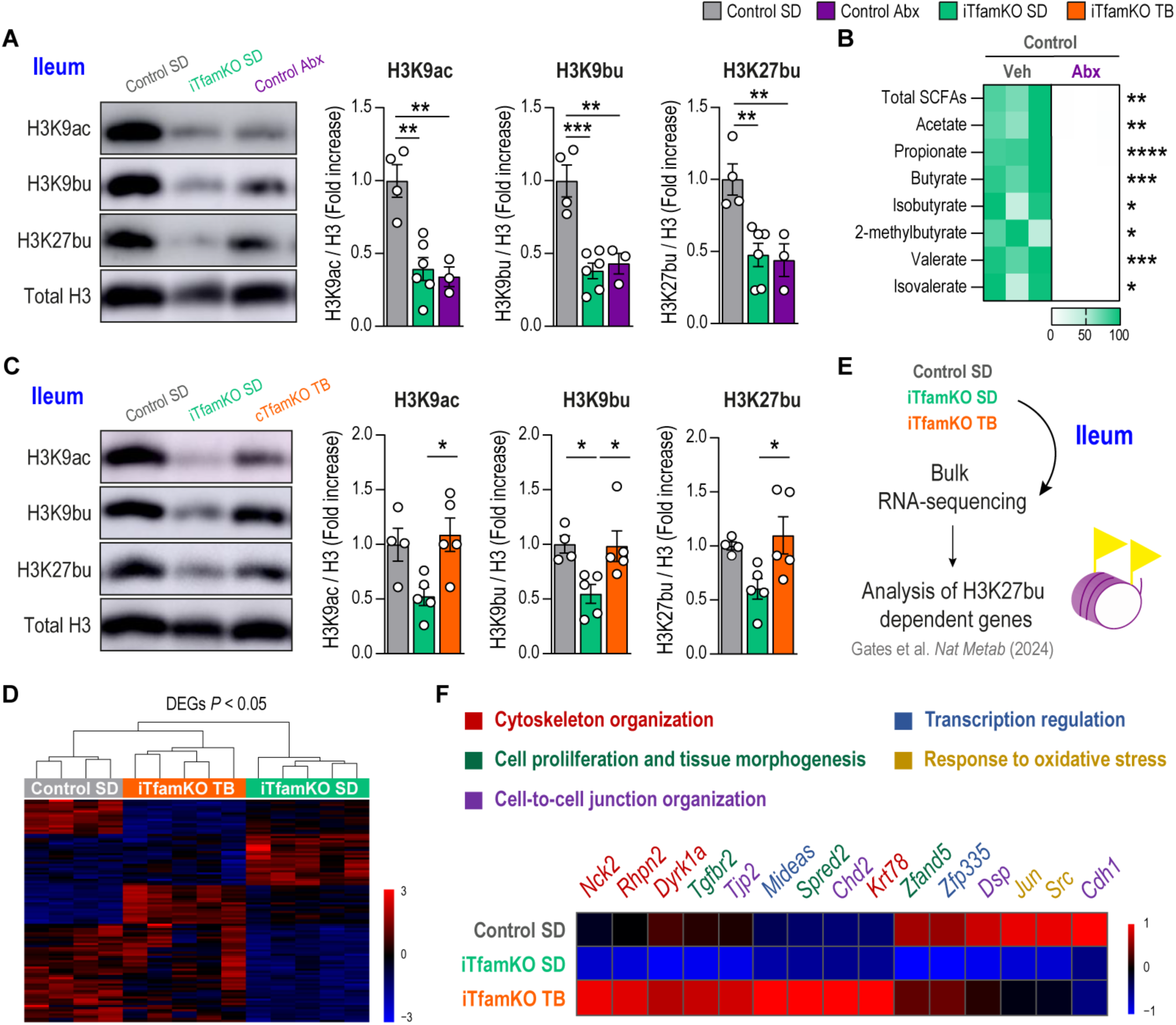
Tributyrin administration restores microbiota dependent epigenetic marks in the small intestine of iTfamKO mice. (**A**) Representative immunoblot and quantification of histone H3 acetylation and butyrylation marks in the small intestine of control, iTfamKO, and antibiotic (Abx)-treated control mice (*n* = 4 to 6). (**B**) Heatmap depicting normalized values of short-chain fatty acid (SCFA) levels in feces of wild-type mice receiving vehicle (Veh) or Abx (*n* = 3). (**C**) Representative immunoblot and quantification of histone H3 acetylation and butyrylation marks in the small intestine of control and iTfamKO mice fed either a standard diet (SD) or a 10% tributyrin-supplemented diet (TB) (*n* = 3 to 6). (**D**) Hierarchical clustering of differentially expressed genes (DEGs) in bulk RNA-sequencing analysis in the ileum. (**E**) Experimental design of the transcriptomic analysis of genes whose expression was previously reported to be dependent on the H3K27bu mark. (**F**) Heatmap depicting expression of H3K27bu-dependent genes in ileum RNA-sequencing data. Data are shown as means ± SEM, where each dot is a biological sample. *P* values were determined by (A and C) one-way analysis of variance (ANOVA) with Tukey’s multiple comparisons test or (B) unpaired Student’s *t* test. \**P* ≤ 0.05; \*\**P* ≤ 0.01; \*\*\**P* ≤ 0.001; and \*\*\*\**P* ≤ 0.0001.

We explored the potential of TB in restoring these histone modifications in iTfamKO mice. Remarkably, TB supplementation restored the levels of H3K9ac, H3K9bu, and H3K27bu in the small intestine of iTfamKO mice when compared with those receiving standard diet (**Figure 7C**). To investigate how TB affected the transcriptomic landscape of the small intestine, we performed bulk RNA-sequencing analysis in the small intestine of control, iTfamKO, and TB-supplemented iTfamKO mice. Hierarchical clustering of differentially expressed genes showed that control and TB-supplemented iTfamKO mice grouped together (**Figure 7D**). This suggested that TB partially restored gene expression alterations in the small intestine of these mice, with upregulation of genes involved in the structural integrity of the mucosa (*Vill*, *Krt80*, *Sh3pxd2b*, *Loxl1*, *Matn2*) and regulation of the immune response (*FoxP3*, *Ctla4*, *Ccr3*, *Ccl22*) (**Figure S8**), which are crucial for the correct performance of the intestinal barrier. Butyrylation of histone H3 has been recently linked to the transcriptional regulation of genes involved in relevant pathways for intestinal homeostasis such as intercellular junctional complexes or oxidative stress^26^. Therefore, we examined the expression of genes previously published to be dependent on the H3K27bu mark^26^ (**Figure 7E**). Focusing on the genes whose expression was repressed in the small intestine of iTfamKO mice compared with controls, we found that TB upregulated genes involved in cell-to-cell junction architecture (*Tjp2*, *Cdh2*, *Dsp*) and cytoskeleton organization (*Nck2*, *Rhpn2*, *Dyrk1a*, *Krt78*), which are relevant for the regulation of the physical barrier of the intestine (**Figures 7F and S9**). Furthermore, we observed increased expression of genes participating in the oxidative stress response (*Jun*, *Src*) as well as in cellular proliferation and tissue morphogenesis, with special emphasis on muscle cell functionality (*Tgfbr2*, *Zfand5*) that may play a role in gut peristalsis (**Figure 7F**). Thus, butyrate supplementation restores epigenetic histone modifications in the intestine of iTfamKO mice correcting the expression of some genes involved in intestinal homeostasis.

Overall, our results indicate that intestinal barrier disruption and gut dysbiosis are hallmarks of mitochondrial dysfunction, leading to reduced SCFA production. Therefore, restoring butyrate levels alleviates systemic organ failure and remarkably extends lifespan in iTfamKO mice.

## Discussion

Disruption of intestinal homeostasis, including increased intestinal permeability and microbial dysbiosis, has been increasingly associated with a wide spectrum of non-communicable pathologies including inflammatory, metabolic, cardiovascular, and neuropsychiatric disorders^30^. Although the precise role of gut dysbiosis in pathology remains scarcely understood, its negative impact on the host has been attributed to its dysregulated metabolic byproducts. SCFAs, key metabolites produced by the gut microbiota through fermentation of dietary and host carbohydrates and proteins, play a central role in safeguarding gut barrier integrity and host physiology. Given its local and systemic beneficial effects, SCFAs have emerged as promising therapeutic agents for a plethora of inflammatory, oncologic, cardiometabolic, and neurological conditions^27^. Thereby, the interplay between host and its commensal microbiota is instrumental to preserve mutualism and ensure health.

In this study, we demonstrate that two distinct mouse models of systemic mitochondrial dysfunction — iTfamKO and mtDNA-mutator mice — exhibit intestinal barrier disruption accompanied by profound bacterial dysbiosis, which is featured by a marked reduction in microbial diversity and loss of commensal symbionts. We observed that mitochondrial failure is associated with gut hyperpermeability in both iTfamKO and mtDNA-mutator mice. Previous reports showed that adult-onset ablation of *Tfam* specifically in the intestinal epithelium impinges on intestinal stem cell (ISC) dynamics blunting enterocyte maturation^37^. mtDNA-mutator mice also show intestinal crypt atrophy and decreased number and activity of proliferative ISCs due to NAD^+^ depletion and subsequent integrated stress response activation^38,39^, which might participate in the breakdown of the gut physical barrier in our mouse models. In addition to the epithelium, *Tfam* depletion also affects smooth muscle cells in the intestinal mucosa of iTfamKO mice, which critically disturb gut peristalsis and intestinal emptying, resulting in signs of constipation as observed in patients living with mitochondrial diseases^20^.

On top of this, mitochondrial metabolism modulates oxygen bioavailability in the intestinal lumen^32^. Therefore, elevated oxygen levels in the intestine of iTfamKO and mtDNA-mutator mice may alter the composition of the commensal microbiota. Importantly, intestinal bacteria show a comparable metabolic profile in iTfamKO and mtDNA-mutator mice characterized by decreased production of SCFAs. These findings reveal a disrupted crosstalk between host mitochondrial function, the intestinal microbiota, and its derived metabolites. This endosymbiotic failure was also shown in mice lacking the mitochondrial heat shock protein 60 specifically in the intestine^33^, which manifest reduced species richness and expansion of disease-associated bacteria prompting intestinal pathology. Beyond intestinal complications^40^, FMT strategies appear to be a feasible tool to alleviate cardiovascular^41^, metabolic^42^, and neurological conditions^43^. In this regard, we have found that transplantation of fecal microbiota from healthy control mice delays signs of disease and extends lifespan in iTfamKO mice, which is associated with restored levels of microbiota-derived SCFAs such as butyrate.

Notably, restoring butyrate mitigates sarcopenia and restores glucose handling metabolism, and kidney function in iTfamKO mice. In line with this, strategies to remodel the intestinal microbiota and its derived SCFAs showed beneficial effects in *Ndusf4^-/-^*mice that mimic Leigh syndrome, a mitochondrial disease that courses with neurological symptoms^44^. Similarly, strategies to restore eubiosis and increase butyrate levels exerted beneficial effects in mouse models of ALS^45,46^, a condition closely linked to mitochondrial dysfunction^47^. Mechanistically, butyrate can modulate post-translational modifications of histones through inhibition of histone deacetylases, and acting as substrate for direct acetylation, propionylation, and butyrylation, linking microbial metabolism to the epigenetic landscape of the host^23,26,48^. Particularly, we observed that the levels of butyrylated histone H3, and expression of several butyryl-H3-dependent genes that are crucial for intestinal homeostasis were decreased in the small intestine of iTfamKO mice. Recovering these epigenetic marks upon TB supplementation was associated with upregulation of genes involved in the physical barrier, and the oxidative stress response in the intestine of these mice. This SCFA-dependent regulation of histone acylations provides a molecular framework by which microbial cues dictate gene expression and host health outcomes. Since the therapeutic effects of tributyrin can be attributed to other molecular mechanisms, future studies should clarify the relevance of butyrate-dependent epigenetic remodeling in disorders showing mitochondrial dysfunction. For example, butyrate can serve as agonist for G-coupled protein receptors and transcription factors, such as Peroxisome Proliferator-Activated Receptors (PPARs)^24,32,48^. Accordingly, PPAR agonists have been shown to boost mitochondrial fitness in a myriad of pathologies presenting with a decline in mitochondrial function^49-52^. In mitochondrially competent cells, butyrate can also act as energy source, which can be metabolized into acetyl-CoA through fatty acid oxidation to help sustain cellular bioenergetics^48^. Upon inducible TFAM depletion, the extent of mitochondrial turnover in the tissues dictates the dilution of pre-existing mtDNA copies and, consequently, the degree of mitochondrial dysfunction. This effect could create a therapeutic window for butyrate to act as an alternative bioenergetic substrate. Butyrate, through its dual role in regulating metabolism and the epigenetic landscape, offers potential therapeutic benefits in various conditions related to mitochondrial dysfunction. Hence, it is tempting to speculate that strategies aiming to restore host-microbiota symbiosis or to normalize microbiota-derived metabolites could ameliorate the progression of diseases associated with deficient mitochondrial function.

## Acknowledgements

We sincerely thank R. Caruso, M. Hasewaga and the Microbiome Core of University of Michigan as well as the Electron Microscopy facility of Centro de Biología Molecular Severo Ochoa (CBM). This research was supported by a European Research Council grant ERC-2021-CoG 101044248-Let T Be (M.M.), Comunidad de Madrid (Spain) grant Y2020/BIO-6350 NutriSION-CM synergy (E.C. and M.M.), Spanish Ministerio de Ciencia e Innovación grant PID2022-141169OB-I00 (M.M.), Comunidad de Madrid (Spain) – Universidad Autónoma de Madrid (UAM) grant SI4/PJI/2024-00166 (E.C.), NIH grant R01 DK095782 (G.N.), Ministerio de Ciencia, Innovación y Universidades (Spain) FPU grants FPU19/02576 (M.M.G.H.), Comunidad de Madrid (Spain) PIPF grant PIPF-2022/SAL-GL-25208 (S.D.-P.), Universidad Autónoma de Madrid FPI-UAM grant (G.S.-H.), and Ministerio de Ciencia, Innovación y Universidades (Spain) Juan de la Cierva-Incorporación grants IJC2018-036850-I (E.G.-R.) and JC2020-044392-I (I.F.-Q.).

## Author contributions

Conceptualization: E.G.-R., M.M.G.H, and M.M. Formal analysis: E.G.-R., M.M.G.H., E.C., P. R.-R., C.S., V.G.-C., N.I., I. B.-L., V. E.-Z., J. O., G.S.-H., E. C., S.D.-P., I. F.-Q., and J. F. A. Funding acquisition: E.C., M.M., and G.N. Investigation: E.G.-R., M.M.G.H., E.C., P. R.-R., C.S., V.G.-C., N.I., I. B.-L., V. E.-Z., J. O., A.F.-A., G.S.-H., E. C., S.D.-P., I. F.-Q., and J. F. A. Resources: R. J.-M., C. G. D., J. A. E., and G.N. Visualization: E.G.-R., M.M.G.H, and M.M. Writing—original draft: E.G.-R., M.M.G.H, and M.M. Writing—review and editing: E.G.-R., M.M.G.H, and M.M.

## Data and materials availability

Microbiota 16*S* rRNA gene sequencing data are publicly available via NCBI with BioProject number PRJNA1291203.

## Materials and methods

### Animal procedures and diets

All animal experimentation procedures were authorized by the Animal Experimentation Ethics Committees of Centro de Biología Molecular Severo Ochoa (CBM) and Centro Superior de Investigaciones Científicas (ProEx 20.5/25) making every effort to minimise mouse discomfort. *Tfam*^fl/fl^*Ubc*^Cre-ERT2^ mice were generated by crossing *Tfam*^fl/fl^ mice with mice expressing the inducible Cre recombinase Cre-ERT2 under the control of the ubiquitin gene (*Ub*^Cre-ERT2^ mice). *Tfam*^fl/fl^*Ubc*^Cre-ERT2^ mouse colony was bred and maintained in the CBM Animal Facility under specific pathogen-free conditions. Both *Tfam*^fl/fl^*Ubc*^Cre-ERT2–/–^ and *Tfam*^fl/fl^*Ubc*^Cre-ERT2+/–^ mice were intraperitoneally administered 20 mg/ml of tamoxifen (Sigma-Aldrich) dissolved in a 10% ethanol, corn oil solution for five consecutive days. For tributyrin supplementation studies, mice were fed either a standard diet (Research Diets, D11112201i) or a 10% (w:w) tributyrin-supplemented standard diet (Research Diets, D24012601i). *PolgA*^D257A/D257^ (*Polg*^mut^) or mtDNA-mutator mice and their controls, *PolgA*^wt^ mice, were bred and maintained in Centro Nacional de Investigaciones Cardiovasculares (CNIC) Animal Facility under specific pathogen-free conditions. Two to five mice were housed per cage separated by genotype and sex and fed ad libitum, receiving cardboard materials as part of the environmental enrichment. Most studies were performed in female mice to facilitate microbiota normalization by mice and/or cage swapping before starting the experiment.

### Glucose tolerance tests

After determination of overnight fasted blood glucose levels, mice were intraperitoneally injected with glucose at a dose of 2 g per kilogram of body weight [10% (w:v)]. Afterwards, blood glucose levels were determined from the blood of mouse tails at 15, 30, 60, 120 and 180 min using Contour next reactive glucose strips and a glucometer (Bayer).

### Body temperature measurements

Body temperature was measured in immobilized mice with their abdomen exposed. An infrared thermometer sensor was then placed below the lower abdomen at approximately 2 to 5 mm away from the abdomen surface while holding the mouse with its body parallel to the ground. Three measurements were performed for each mouse.

### Forelimb grip strength assessment

Mice were held by their tails and forelimb grip strength was measured as tension force using a digital force transducer (Grip Strength Meter, Bioseb). Five to seven measurements per trial were performed for each mouse, with a few seconds resting period between measurements.

### Rotarod test

Motor coordination was assessed in an accelerating rotarod apparatus (Ugo Basile). Mice were trained for two consecutive days at constant speed: on the first day, four times at 4 revolutions per minute (rpm) for 1 min and, on the second day, four times at 8 rpm for 1 min. The day afterwards, the rotarod instrument was set to progressively accelerate from 4 to 40 rpm for 5 min. During the accelerating trials, the latency to fall from the rod was measured. Mice were tested four times.

### Open field test

For behavioral experiments, mice were simultaneously transferred to a behavior room where they habituated for several days before the start of the tests. The light/dark cycle was 12/12 hours (lights on at 8:00 am). All the experiments were performed during the light phase. To minimize variability, each animal was always tested at the same time of the day. Open field test was measured in a 43 × 43 cm square test arena and recorded for 3 days. Mice were first habituated to the dark for 30 min and then placed in the illuminated arena for 10 min. Free movement was recorded and analyzed using ANY-maze™ Video Tracking System software (Stoelting Co.). To assess locomotion, total distance travelled, exploration speed, and time immobile were quantified.

### Nest building test

Nest building performance was performed as formerly described^53^. Briefly, mice were individualized overnight in clean cages containing bedding but no environmental enrichment. Then, an intact new nestlet was placed inside the cage. Next morning, the nest building was scored as following: nestlet mostly untouched (1 point); nestlet partially torn but mainly flat (2 points); nestlet mostly torn but no nest identifiable (3 points); identifiable but mainly flat nest (4 points); perfectly built nest with walls covering the mouse body (5 points).

### Hematological analyses

Blood was extracted by submandibular vein puncture in Microvette^®^ EDTA K2 tubes. Supernatant plasma was collected from anticoagulated samples after centrifugation at 10000*g* for 12 min at 4°C. Finally, plasma was assessed with an Abacus Junior or Element HT5 hematology analyzer.

### Intestinal permeability assay

Mice fasted 2 hours were orally gavaged with a 1:3-1:4 (v:v) dilution of 250 mg/ml 4-kDa fluorescein isothiocyanate (FITC)-dextran probe (Sigma-Aldrich) at a dose of 0.6 g per kilogram of body weight. After 2 hours, 100 to 120 μl of blood was collected in BD Microtainer^®^ tubes from the facial vein of mice and centrifuged at 6000*g* for 10 min at 4 °C to obtain the serum fraction. FITC-dextran measurements were performed in duplicates by fluorimetric quantification of mouse serum mixed with an equal volume of phosphate buffered saline (PBS) [1:9 (v:v)]. Dilutions of non-treated mouse serum in FITC-dextran diluted with PBS were used as a standard curve to calculate blood FITC-dextran concentrations. One hundred microliters of standards or diluted sera was measured in a FLUOstar OPTIMA® (BMG Labtech) 96-plate reader at an excitation wavelength of 492 nm and an emission wavelength of 525 nm.

### Microbiota depletion experiments

Mice were randomly assigned to vehicle or antibiotics (Abx) treatment groups. Mice in the latter were administered a cocktail of neomycin (1 mg/ml) (Nzytech), ampicillin (1 mg/ml) (Nzytech), metronidazole (1 mg/ml) (Sigma-Aldrich) and vancomycin (0.5 mg/ml) (Sigma-Aldrich) in autoclaved drinking water supplemented with sucrose (2 mg/ml) (2 mg/ml) to improve palatability and renewed weekly. Vehicle mice were treated with sucrose water without antibiotics for the same time period.

### Fecal microbiota transplantation (FMT)

FMTs were performed as formerly described^54^. In brief, mice fasted 6 hours were orally gavaged for 3 consecutive days with 200 μl of an antibiotic cocktail consisting of neomycin (1 mg/ml) (Nzytech), ampicillin (1 mg/ml) (Nzytech), metronidazole (1 mg/ml) (Sigma-Aldrich) and vancomycin (0.5 mg/ml) (Sigma-Aldrich) in autoclaved water. The day afterwards, four to five fresh fecal pellets were pooled from donor mice in 600 μl of reduced buffer (0.5 mg/ml cysteine and 0.2 mg/ml Na_2_S in PBS) and vortexed for 1 min. Homogenates were then centrifuged at 500*g* for 5 min to remove large particles. Finally, 200 μl of fecal slurry was orally gavaged to recipient mice fasted for 4 hours twice a week for 2 weeks and then, once a week until sacrifice. Following FMT, the remaining slurry was applied on the fur of recipient mice, and their cages were replenished with fresh fecal pellets and dirty bedding from donor mice to ensure coprophagia.

### Histological, immunohistochemical, and immunofluorescence analysis

#### Histological analysis

Organs were collected from euthanized mice after intracardiac perfusion with cold PBS. In particular, small intestine and colon were harvested by cutting below the stomach and above the cecum, and just below the cecum until the rectum, respectively. Fat was removed, and lumen was flushed with cold PBS to expel fecal and mucus content. Organs were fixed in 10% neutral buffered formalin for 24 hours and dehydrated in 70% ethanol until processing. Dehydrated organs were embedded in paraffin and further processed with a microtome to obtain sections. Afterwards, sections were deparaffinized and stained as detailed in figure legends for their histological examination. Images were captured using the 5X objective of a vertical microscope AxioImager M1 (Zeiss) connected to a DMC6200 camera (Leica) and the LasX software (v4.13). Crypt depth and villus height were measured from the bottom of the crypt to the crypt-villus junction, and from there to the tip of the villus, respectively. Twenty-five to fifty crypt–villus units per mouse were quantified in a single-blind manner. Elastic lamina breaks, defined as interruptions in the elastic fibers, were counted in the entire medial layer of 3 consecutive cross-sections per mouse and the mean number of breaks was calculated. Image quantification was performed using the NanoZoomer Digital Pathology software (v2.7.25).

#### Immunofluorescence analysis

For brain immunofluorescence staining, the left hemisphere of each brain was fixed by immersion in 4% paraformaldehyde (PFA) in PBS for 24 hours at room temperature (RT) and stored in PBS at 4°C. Then, fixed hemispheres were embedded in a solution of 10% sucrose–4% agarose in phosphate buffer (PB). The resulting blocks were cut with a vibratome (Leica) obtaining 40 μm-thick sagittal sections which were stored at – 20°C in 40% glycerol, 30% ethylene glycol in 0.2 M PB, pH 7.4. Brain slices were washed with Tris buffered saline (TBS), permeabilized and blocked with blocking solution (0.5% Triton X–100 and 2% BSA in TBS) for 1 hour at RT. Sections were then incubated overnight with mouse anti-NeuN (1:500 dilution; Millipore, MAB377) diluted in blocking solution at 4°C. After 3 washes with TBS, each section was incubated with secondary antibodies conjugated to fluorophores 1 hour at RT. They were washed again with TBS and nuclei were stained with DAPI for 5 min (1/5000 dilution; Merck, 268298). Finally, sections were mounted on slides using Prolong Glass Mounting Medium (Invitrogen). Images were acquired using the oil immersion 25X objective of a LSM710 confocal microscope coupled with a vertical microscope AxioImager M2 (Zeiss) and the ZEN Black 2010 software and were analyzed with the ImageJ software (v1.53n). Lipofuscin granules were quantified in a single-blind manner in brain slides owing to their auto-fluorescent properties.

#### Electron microscopy

After collection, a portion of the small intestine and the colon were fixed with 4% PFA 2% glutaraldehyde in 0.1 M PB, pH 7.4. Tissues were post-fixed with 1% osmiun tetroxide and 1% potassium ferrocyanide in water for 1 hour at 4°C, dehydrated with lowering concentrations of ethanol and finally embedded in resin EPON. 80 nm-sections of the embedded tissue were obtained using an Ultracult E ultramicrotome and mounted on carbon-coated copper slot grids. Sections were counterstained with 2% uranyl acetate and lead citrate for 1 hour. Preparations were examined with a transmission electron microscope (JEM1400 Flash, Jeol) and images were acquired with a CMOS Oneview camera (Gatan). The area of mitochondria and the number of body inclusions per mitochondria were quantified in a single-blind manner using the ImageJ software (v1.53n).

#### Immunohistochemical analysis

For immunohistochemical staining, deparaffinized small intestine sections were rehydrated and boiled in citrate buffer (10 mM citrate buffer, 0.05%Triton X-100, pH 6) to retrieve antigens. Then, sections were blocked in 10% goat serum, 5% horse serum, 0.05% TritonX-100, and 2% BSA in PBS for 45 min. Endogenous peroxidase and biotin were blocked with 1% hydrogen peroxide-methanol for 10 min and a biotin blocking kit (Vector Laboratories), respectively. Sections were incubated with a goat anti-Ki67 antibody (RD Systems, AF3667) and color was obtained with 3,3’-diaminobenzidine (Vector Laboratories). Sections were counterstained with hematoxylin and mounted in DPX (Fluka). Slides were scanned with a NanoZoomer-RS scanner (Hamamatsu). The number of Ki67-positive cells was quantified in a single-blind manner using NanoZoomer Digital Pathology software (v2.7.25).

### Immunoblot

For Western blot analysis, frozen tissues were dissected and homogenized in RIPA buffer (20 mM Tris–HCl, pH 7.5, 150 mM NaCl, 1 mM EDTA, 1 mM EGTA, 1% NP−40, 1% sodium deoxycholate, 0.1% SDS) with proteases (cOmplete^TM^, Sigma-Aldrich) and phosphatases inhibitors (Sigma-Aldrich). The concentration of proteins in homogenized samples was determined using the Pierce^TM^ BCA Protein Assay Kit (Thermo Fisher Scientific) or the DC Protein Assay (Biorad), following manufacturer’s instructions. Thirty to 50 μg of brain lysates, or 20 μg of small intestine lysates were prepared in Laemmli buffer (25 mM Tris–HCl pH 6.8, 1% SDS, 3.5% glycerol, 0.4% 2-mercaptoethanol and 0.04% bromophenol blue) and separated by electrophoresis in polyacrylamide gels in the presence of sodium dodecylsulfate (SDS) at constant voltage.

Proteins were then transferred to nitrocellulose membranes by wet transfer for brain lysates or using the Trans-Blot® Turbo^TM^ system (Biorad) for small intestine lysates, and after blocking with 5% BSA or 5% milk in 0.1% Tween-20 in TBS [T-TBS], membranes were incubated overnight with rabbit anti-TFAM (1:1000 dilution; Proteintech, 22586), rabbit anti-calnexin (1:10000 dilution; Abcam, ab22595), rabbit anti-tau-P Ser 396 (1:1000 dilution; Life Technologies, 44752G), mouse anti-PSD95 (1:1000 dilution; BD Biosciences, 610495) or mouse anti-vinculin (1:10000 dilution; Abcam, ab129002) for brain lysates, or rabbit anti-H3K9ac (1:5000 dilution; Abcam, ab4441), rabbit anti-H3K9bu (1:500 dilution; PTMBio, PTM-305), rabbit anti-H3K27bu (1:1000 dilution; Merck, ABE2854), or rabbit anti-H3 (1:1000 dilution; Abcam, ab1791) for small intestine lysates diluted in T-TBS at 4°C with gentle agitation. After washing with T-TBS, membranes were incubated with the corresponding secondary antibodies coupled to horseradish peroxidase at 1:2500 to 1:15000 dilution for 1 hour at RT. Protein signal was detected with Pierce^TM^ ECL Western Blotting Substrate (Thermo Fisher Scientific) and chemiluminescence was measured with an Amersham™ Imager 680 (GE Healthcare). Each protein of interest was quantified using the ImageJ software (v1.53n), by measuring the average pixel intensity of the corresponding band and normalizing the value to the respective control protein.

### RNA Isolation and qPCR Analysis

Total RNA extraction was performed with a MagNA Lyser® homogenizer (Roche) using 1 cm of frozen tissue in 700 μl of TRIzol® reagent (Invitrogen). The RNA from the resulting aqueous phase was further purified using the RNeasy Mini Kit (QIAGEN) following manufacturer’s instructions. Quality and quantity were determined by measuring the absorbance at 260 and 280 nm on a Nanodrop One spectrophotometer (Thermo Fisher Scientific).

cDNA libraries were prepared from total RNA and then, validated and quantified by an Agilent 4200 TapeStation in Haplox. After passing library inspection, stranded mRNA was sequenced either on the Illumina Novaseq Xplus platform, and FastQ files were generated containing nucleotide data and quality scores for each position. The quality of FastQ files was checked using FastQC (v0.11.9) RNA-sequencing reads were mapped to the *Mus musculus* reference genome GRCm39 using either Hisat2 (v2.2.1) or STAR (v2.5.2) software. Reads were then pre-processed with SAMtools (v1.13) to transform Sequence Alignment/Map files into Binary Alignment/Map files and sorted. The number of reads covered by each gene was calculated by HTSeq-Count (v1.99.2).

Downstream data analysis was performed with R (v4.3.2). Differential gene expression (DEG) analysis was performed using DESeq2 (v1.44). Genes with *P* value < 0.05 and |log_2_-fold change| > 0.6 were determined to show statistically significant differences in group comparison. Over-representation analysis (ORA) and gene set enrichment analysis (GSEA) were performed using clusterProfiler (v4.8.3) package in GO, KEGG, WikiPathways, Reactome and the Hallmarks of the Molecular signatures databases. PCA plots, chord diagrams and heatmaps were visualized by using ggplot2 (v3.4.4), circlize (v0.4.15) and pheatmap (v1.0.12), respectively.

For qPCR analysis, reverse transcription was performed using 500 ng of RNA extracts and the Maxima™ First Strand cDNA Synthesis Kit and dsDNase (Thermo Fisher Scientific). The reaction was performed in a ProFlex™ PCR System thermocycler (Applied Biosystems) at 25°C for 10 min, followed by 50°C for 15 min, and 85°C for 5 min. cDNA samples were then cooled to 4°C and stored at − 20°C for qPCR analysis. Amplification conditions were determined by the primers to present amplification efficiency close to 100% and a single peak in melt-curve analyses. We included 1:15-1:25 dilution of cDNA samples in the qPCR reaction, along with 5 μl of polymerase (Go Taq® Master Mix, Promega) and 0.5 μl of each forward and reverse primer solution (5 μM stock) added to 384-well plates. The reaction was run on a Bio-Rad CFX 384 thermocycle. The following 5’- 3’ primer pairs (FW, RV. Sigma-Aldrich) were used in this study:

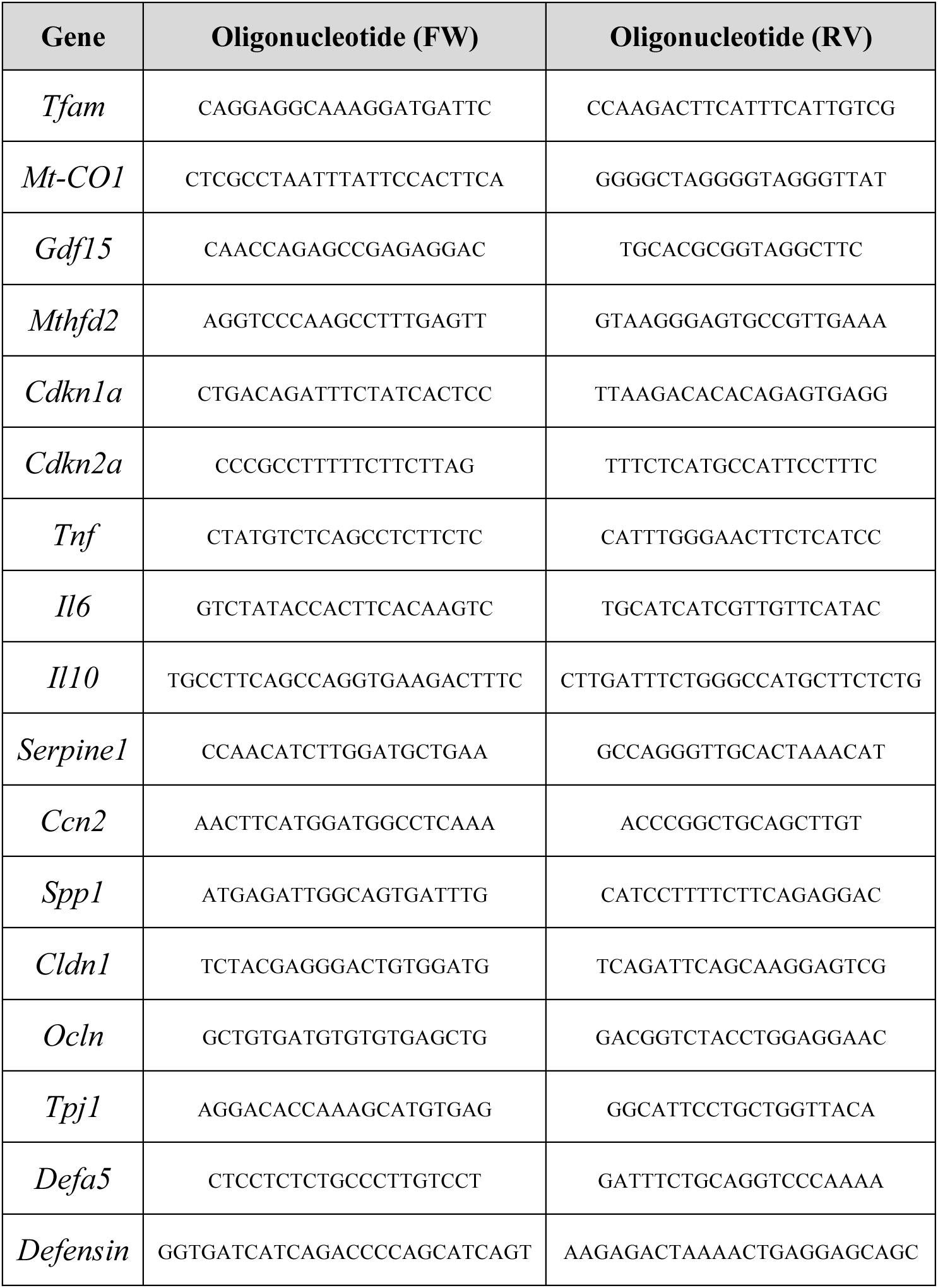
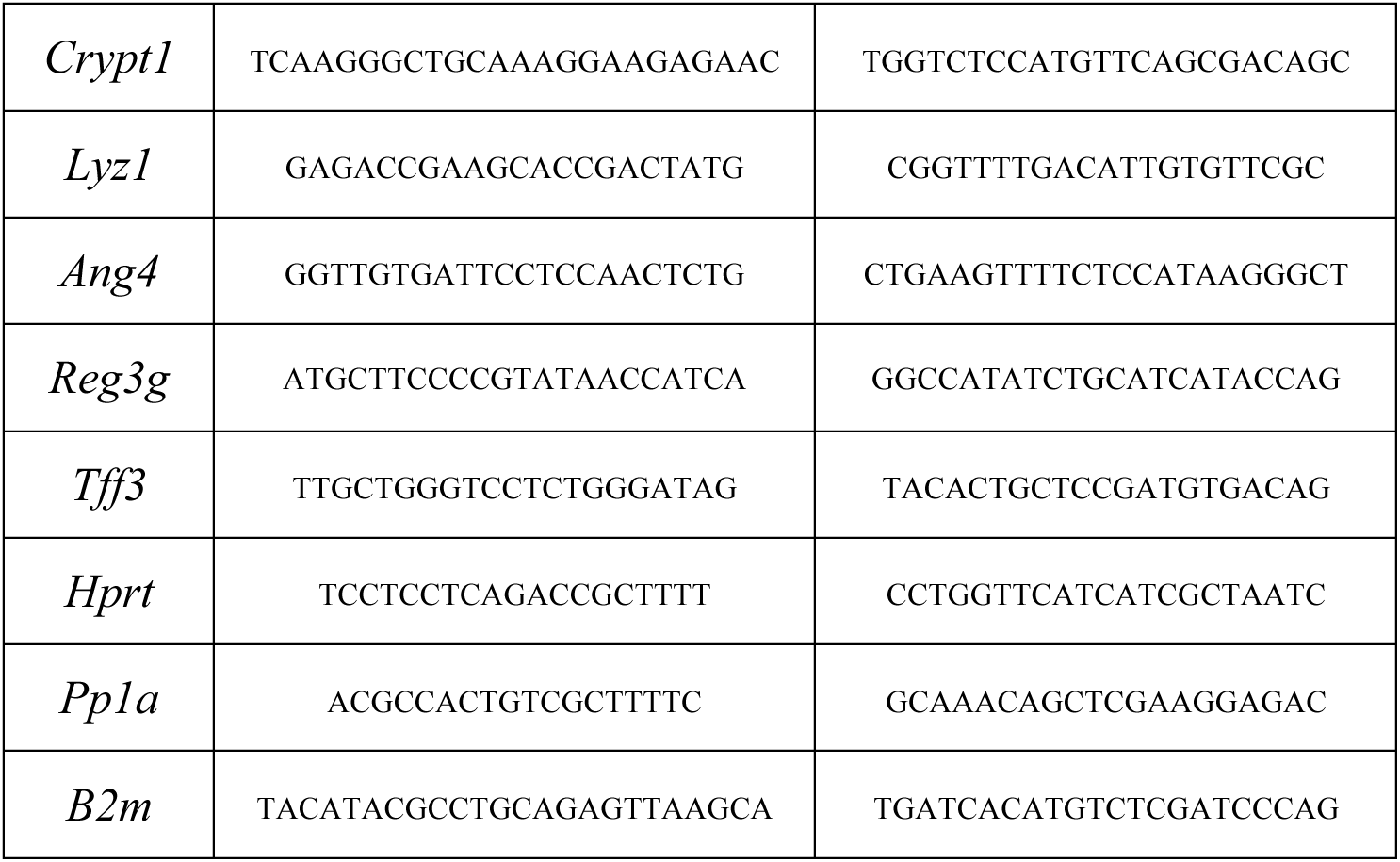

Quantitative RT-PCR data were analyzed using the 2^-ΔΔCt^ method to calculate the relative changes in gene expression, where ΔΔCt is the difference between the problem sample Ct values, and the control ones. The relative expression in each figure represents the induction levels of the gene of interest relative to *Hprt*, *Pp1a* or *B2m*.

### Urine protein analysis

Urine was collected from mice after immobilizing them on top of a piece of Parafilm and immediately frozen at – 80°C. Ten microlitres of urine samples or 0.05 mg/ml BSA (Biorad) in Laemmli buffer were loaded into a 12% polyacrylamide gels in the presence of SDS at constant voltage to perform electrophoresis. Electrophoresis gel was then stained with a 0.1% Coomassie blue (w:v) in 20% methanol and 10% acetic acid solution by heating the gel for 1 min in a microwave and placing it on a shaker for 30 min at RT. Afterwards, gel was destained using a 20% methanol 10% acetic acid solution by heating it for 30s in a microwave and, then, wash it several times with the indicated solution until bands start to be visible. Images were acquired with an Amersham™ Imager 680 (GE Healthcare). The band corresponding to albumin was quantified using the ImageJ software (v1.53n).

### Luminex detection of cytokines

Blood (100-120 μl) was collected in BD Microtainer^®^ tubes from the facial vein or after culling mice and centrifuged at 6000*g* for 10 min at 4°C to obtain the serum fraction. Cytokines from serum samples were quantified using the multiplexed bead-based immunoassay Cytokine & Chemokine 26-Plex Mouse ProcartaPlex™ Panel 1 (Invitrogen, EPX260-26088-901) following manufacturer’s instructions.

### LPS-binding protein (LBP) ELISA quantification

Serum samples from mice were diluted and processed following manufacturer’s instructions (Enzyme Immunoassay for quantification of mouse LBP, Biometec). The plate was measured in a Dynex Opsys MR^TM^ 96-plate reader (Aspect Scientific) at an excitation wavelength of 450 nm and an emission wavelength of 630 nm.

### Fecal output and water content analyses in feces

Mice were individualized in clean cages without bedding or environmental enrichment for 30 min, and the number of feces per cage was enumerated at the end of the experiment. A fresh fecal sample from each animal was immediately weighed and dried at 50°C overnight. Afterward, samples were weighed again, and the difference was considered the fecal water content and expressed as a percentage.

### Microbiota analysis by *16S* rRNA gene sequencing

*16S* rRNA gene sequence analyses were performed as formerly reported^55^. In brief, bacterial DNA from ileal, colonic, and fecal samples was extracted using the E.Z.N.A stool DNA kit (Omega Biotek) following manufacturer’s instructions. Amplicons of the V4 region of the *16S* rRNA gene were obtained in each sample using the following 5′-3′ primer pair:

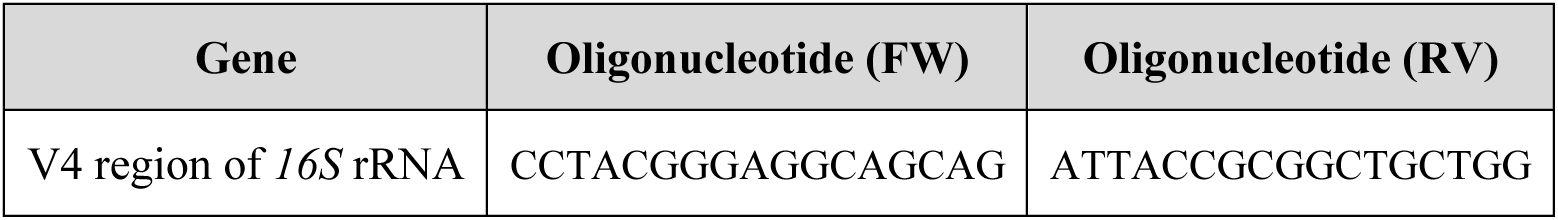

Libraries were then sequenced using an Illumina MiSeq instrument, and sequences were curated and analyzed using the mothur (v.1.40.5) software package.

### Short-chain fatty acid (SCFA) quantification by UPLC-MS/MS

Standards of eight straight and branched-chain SCFAs [acetic acid (AA), propionic acid (PA), isobutyric acid (i-BA), butyric acid (BA), 2-methylbutyric acid (2-Me-BA), isovaleric acid (i-VA), valeric acid (VA), and 3-methylvaleric acid (3-Me-VA)], 3-nitrophenylhydrazine (3NPH), N-(3-dimethylaminopropyl)-N′-ethylcarbodiimide hydrochloride (EDC), formic acid, and pyridine were purchased from Sigma-Aldrich. Acetonitrile (ACN) was purchased from VWR. Calibration curves were constructed using a mixed SCFA solution in ACN-water (50:50, v/v), with ranges of 1 to 10,000 µM for AA and 0.1 to 1000 µM for PA, i-BA, BA, 2-Me-BA, i-VA, and VA.

For fecal samples, SCFAs were extracted from 50 mg of feces using 100 µl of ACN-water (50:50, v/v) containing 5 µM internal standard, homogenized with a FastPrep-24 5G system (MP Biomedicals), incubated at 800 rpm for 15 min at 10°C, and centrifuged at 14,000*g* for 30 min at 4°C. For serum samples, SCFAs were extracted by adding 60 μl of ice-cold methanol (containing IS) to 30 μl of serum, following incubation on ice for 1h. Then, the suspension was centrifuged at 14000*g* for 15 min and 4 °C. Forty microliters of the supernatants were derivatized by mixing with 20 µl of 200 mM 3NPH and 120 mM EDC/6% pyridine in ACN-water (50:50, v:v) and incubated for 30 min at 40°C. After cooling to RT, fecal and serum samples were diluted 20-fold or 2-fold, respectively, with 10% ACN in water, centrifuged at 14,000*g* for 10 min at 4°C, and analyzed by UPLC-MS/MS. Standards and blanks were processed using the same derivatization protocol.

SCFA analysis was conducted following a previously described protocol^56^. Analyses were carried out using an Agilent 1260 Infinity II system coupled to an Ultivo 6465 Triple Quadrupole LC-MS, equipped with an Agilent Jet Stream ESI source and controlled via MassHunter Workstation (Agilent Technologies). Multiple Reaction Monitor (MRM) conditions acquisition modes were optimized by direct infusion of the 3NPH derivatives as follows:

| **3NPH-SCFA**<br>**derivatives** | **Precursor ion**<br>**(m/z)** | **Product ion**<br>**(m/z)** | **Fragmentor**<br>**voltage (V)** | **Collision**<br>**energy (eV)** |
| --- | --- | --- | --- | --- |
| AA (C2) | 194.1 | **137**<br>46 | 92 | **16**<br>44 |
| PA (C3) | 208.1 | **137**<br>46 | 92 | **20**<br>48 |
| i-BA/BA (C4) | 222.1 | **137**<br>107 | 92 | **20**<br>28 |
| 2-Me-BA/i-VA/VA (C5) | 236.1 | **137**<br>107 | 102 | **20**<br>32 |
| 2-Me-VA (C6) | 250.1 | **137**<br>107 | 102 | **20**<br>31 |

### Statistical analyses and figure design

Statistical analyses were performed using GraphPad Prism 9 or Past 3.22 software. Outliers were identified and excluded by the ROUT method (5%). If data followed a normal distribution after applying the Shapiro–Wilk test, comparisons between two datasets were performed using the unpaired two-tailed Student’s *t* test, and comparisons between more than two datasets were performed using the one or two-way analysis of variance (ANOVA) or mixed-effects analysis with Tukey’s or Šidák’s multiple comparison tests. If not, comparisons between two or more datasets were calculated using the non-parametric Mann–Whitney *U* test and Kruskal–Wallis *H* test with Dunn’s multiple comparison test, respectively. For categorical variables, Fisher’s exact test was used. For survival curves, the log-rank Mantel–Cox test was applied. Permutational multivariate analysis of variance (PERMANOVA) was performed based on 9999 permutations. Differences with *P* values ≤ 0.05 were considered significant. \**P* ≤ 0.05; *\**\**P* ≤ 0.01; *\*\**\**P* ≤ 0.001; and \*\*\*\**P* ≤ 0.0001. Unless otherwise stated, experimental data were represented as means ± the standard error of the mean (SEM), where each dot was an individual biological sample of each experimental group. In the figure legends, (*n*) denotes the number of pooled mice per group in the experiment. Box-and-whisker plots represent the interquartile range between the first and third quartiles (25th and 75th percentiles, respectively), the median, and the maximal and minimal values. Violin plots represent the interquartile range between the first and third quartiles (25th and 75th percentiles, respectively) and the median. Figures were designed using GraphPad Prism 9 and Adobe Illustrator (v29.0.1).

## Figure legends

**Figure S1.**
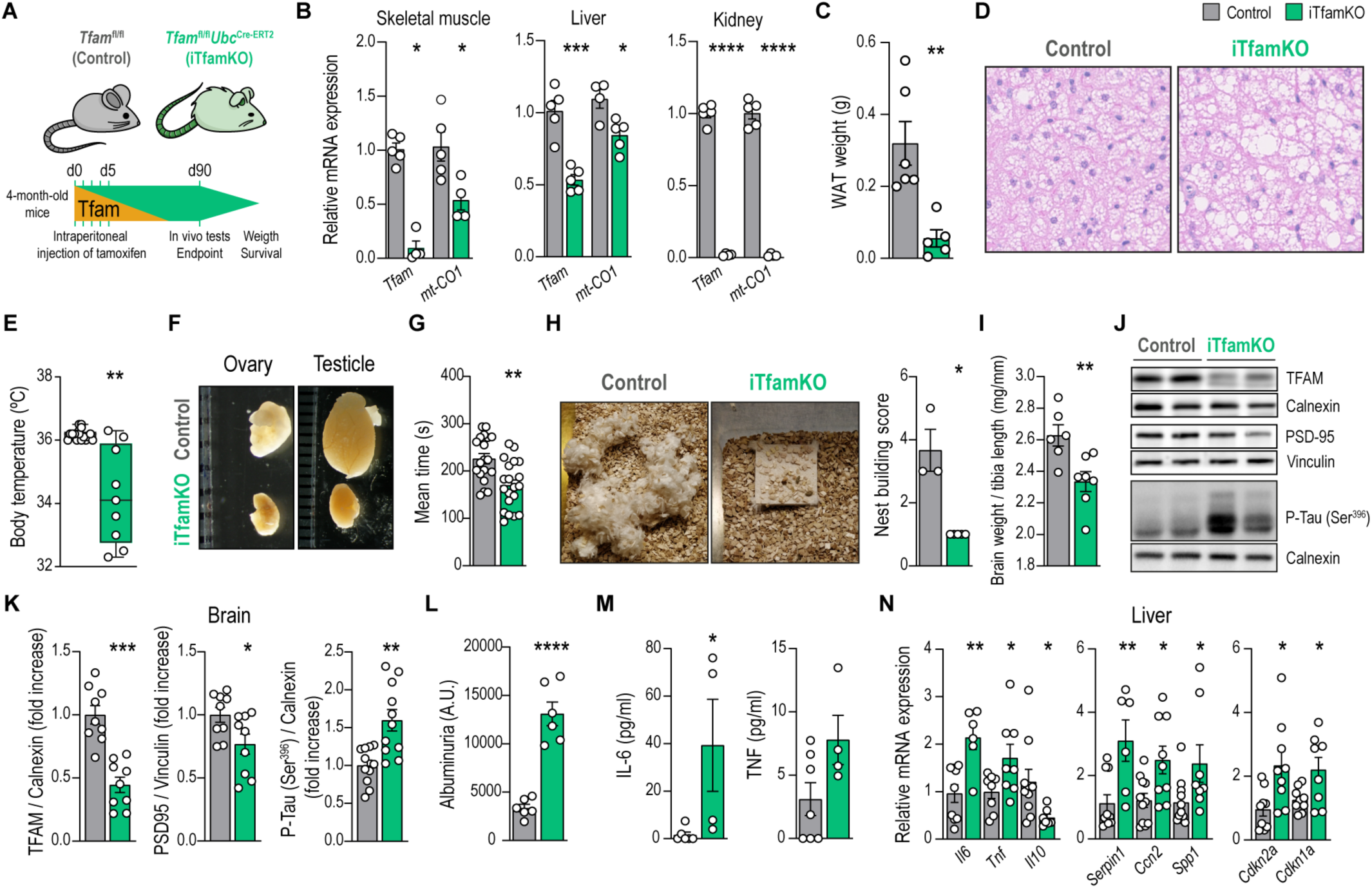
Inducible deletion of *Tfam* in adult mice precipitates multimorbidity. (**A**) Experimental design of tamoxifen inducible deletion of *Tfam* in adult mice. (**B**) Relative mRNA levels of *Tfam* in the skeletal muscle, liver, and kidney of control and cTfamKO mice 90 days after tamoxifen administration (*n* = 4 to 5). (**C**) Quantification of gonadal white adipose tissue (WAT) weight (*n* = 5 to 6). (**D**) Representative hematoxylin and eosin–stained sections of brown adipose tissue. (**E**) Quantification of mouse body temperature (*n* = 9 to 16). (**F**) Representative picture of mouse gonads. (**G**) Rotarod test performance expressed as the mean time spent on the rotating rod in all trials combined (*n* = 19 to 20). (**H**) Representative picture of a mouse nest and its respective score (*n* = 3 to 5). (**I**) Quantification of brain weight normalized to tibia length (*n* = 6 to 7). (**J**) Representative immunoblot of TFAM, PSD95, and phospho-Tau (Ser396) proteins in brain samples. (**K**) Densitometry analysis of brain immunoblot (*n* = 8 to 12). (**L**) Quantification of albuminuria (*n* = 6). (**M**) Concentration of IL-6 and TNF in the serum (*n* = 4 to 7). (**N**) Relative mRNA levels of genes encoding inflammatory mediators (*Il6*, *Tnf*, *Il10*), pro-fibrotic factors (*Serpin1*, *Ccn2*, *Spp1*), and senescence-associated markers (*Cdkn2a*, *Cdkn1a*) in the liver (*n* = 6 to 11). Data are shown as means ± SEM, where each dot is a biological sample. *P* values were determined by (B, C, G, I, and K to N) unpaired Student’s *t* test, (E) Wilcoxon text or (H) two-tailed Mann-Whitney *U* test. \**P* ≤ 0.05; \*\**P* ≤ 0.01; \*\*\**P* ≤ 0.001; and \*\*\*\**P* ≤ 0.0001.

**Figure S2.**
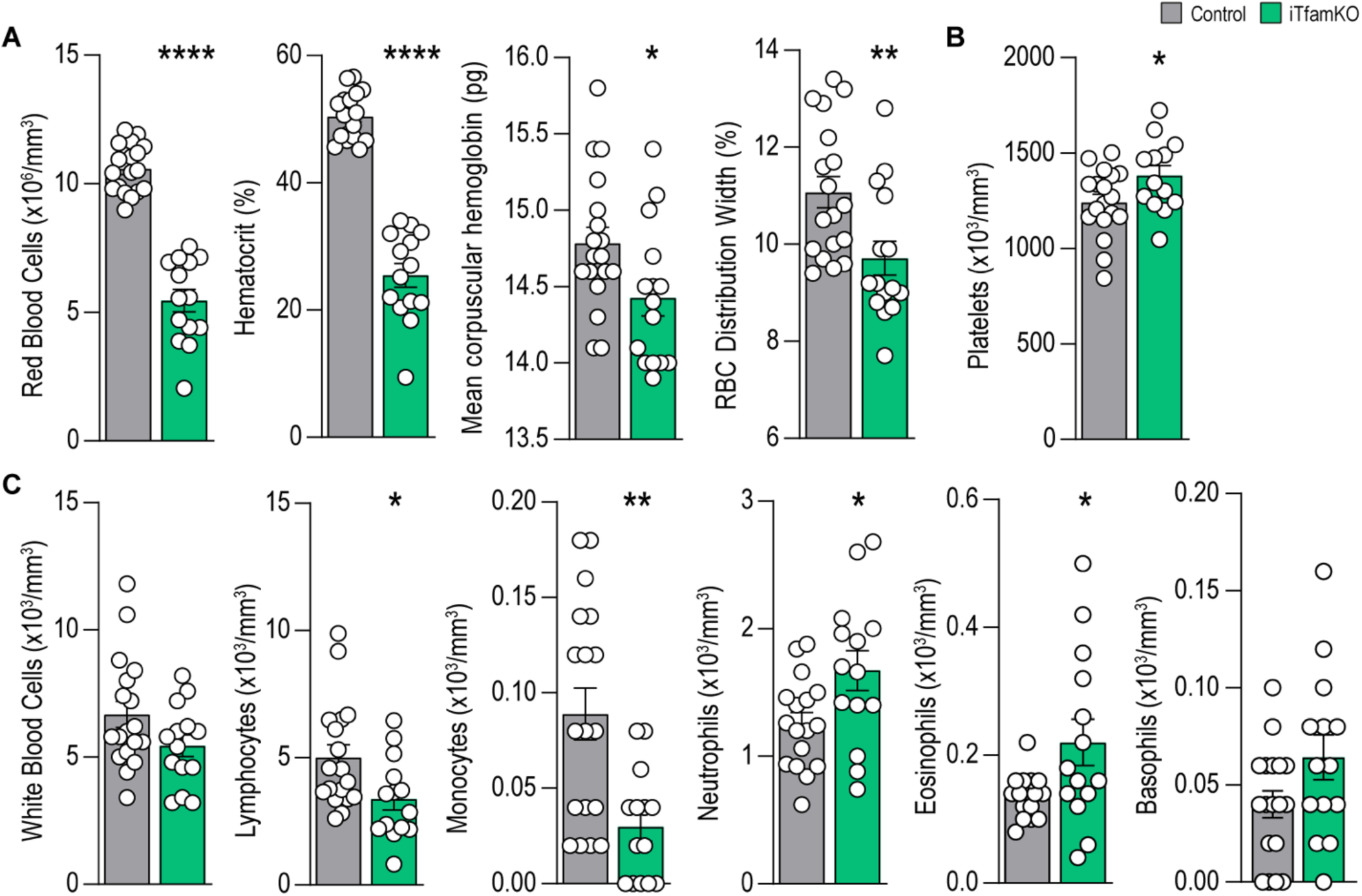
iTfamKO mice exhibit severe alterations in the hematopoietic cell compartment. (**A**) Quantification of hematologic parameters related to red blood cell (RBC) biology in control and iTfamKO mice 90 days after tamoxifen administration (*n* = 14 to 18). (**B**) Quantification of blood platelets (*n* = 13 to 18). (**C**) Quantification of hematologic parameters related to white blood cell biology (*n* = 14 to 18). Data are shown as means ± SEM, where each dot is a biological sample. *P* values were determined by unpaired Student’s *t* test. \**P* ≤ 0.05; \*\**P* ≤ 0.01; and \*\*\*\**P* ≤ 0.0001.

**Figure S3.**
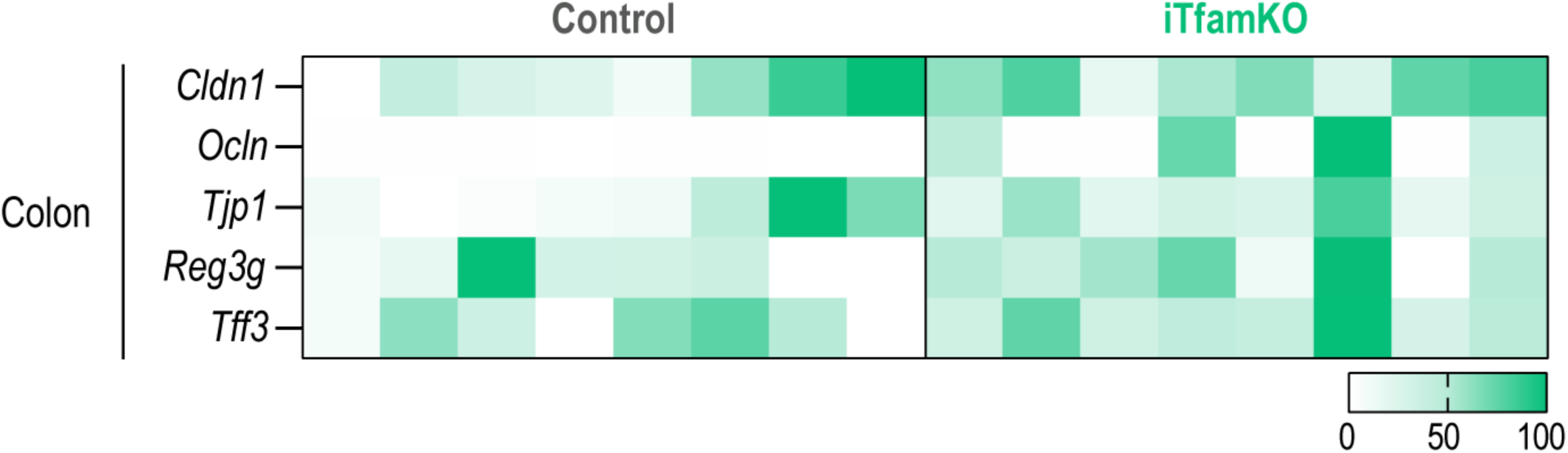
Analysis of barrier integrity genes in the colon of iTfamKO mice. Heatmap depicting normalized values of qPCR analysis of genes associated with tight junctions (*Cldn1*, *Ocln*, *Tjp1*), antimicrobial peptides (*Reg3g*), and barrier function (*Tff3*) in the colon of control and iTfamKO mice (*n* = 7 to 9). *P* values were determined by unpaired Student’s *t* test.

**Figure S4.**
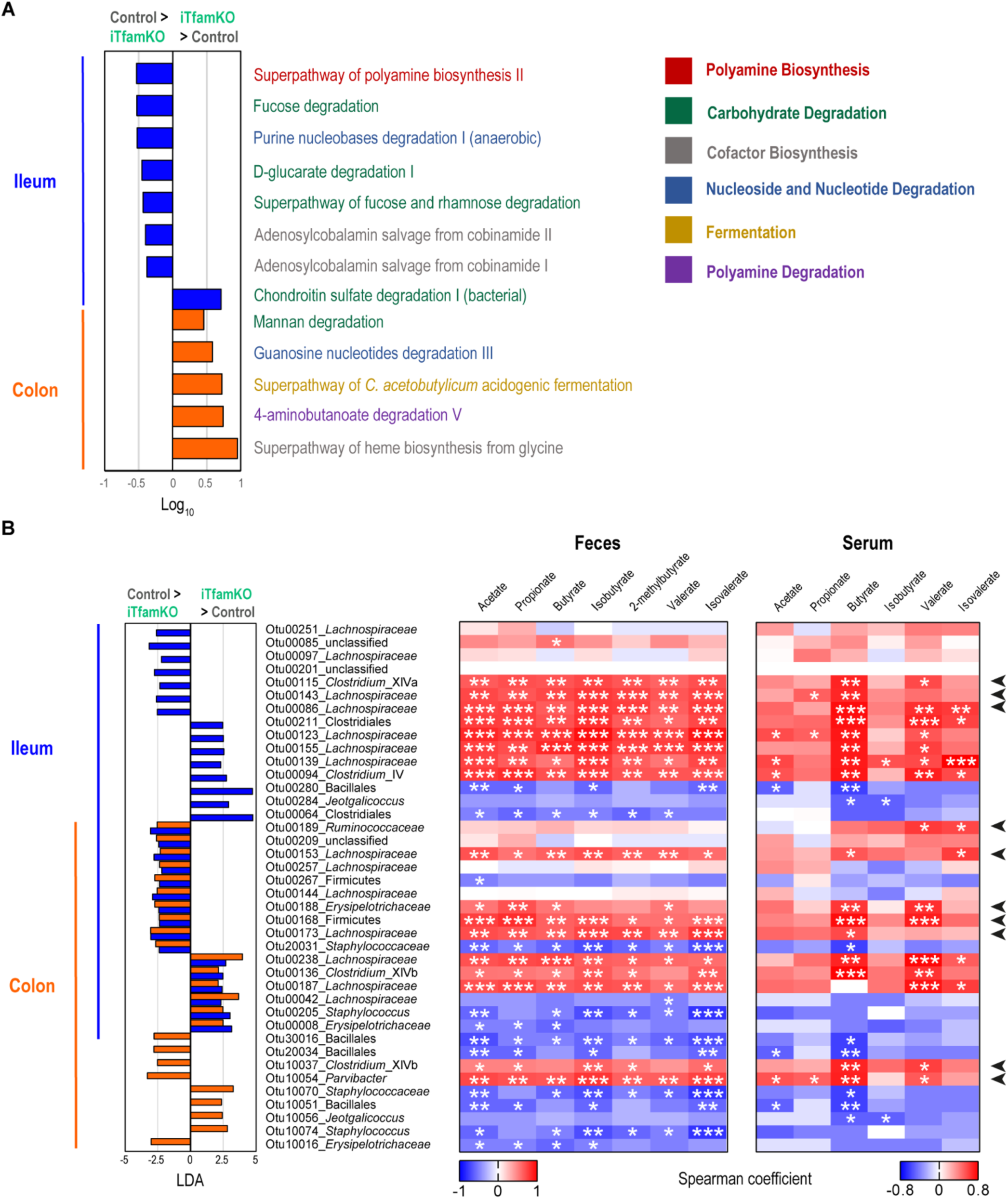
Predictive metabolic profiling of the gut microbiota in iTfamKO mice. (**A**) PICRUSt2 prediction of metabolic pathways in ileal and colonic metagenomic data of control and iTfamKO mice (FDR, *Q* < 0.05; |SNR| > 0.5; |LDA| > -1; |fold change| > 2; maximal index > 1000). (**b**) Left: differentially abundant operational taxonomic units (OTUs) depicted with lineal discriminant analysis (LDA) values of linear discriminant effect size (LEfSe, *P* < 0.05; FDR, *Q* < 0.05; |SNR| > 0.5; |LDA| > 2; |fold change| > 10; maximal abundance > 0.001) comparing the ileum and colonic microbiota in control versus iTfamKO mice. Right: heatmap depicting Spearman’s rank correlation coefficients between differentially abundant OTUs in ileal and colon-resident microbiota, and the concentration of SCFAs in the feces and the serum of mice. *P* values were determined by PICRUSt2 software. \**P* ≤ 0.05; \*\**P* ≤ 0.01; and \*\*\**P* ≤ 0.001.

**Figure S5.**
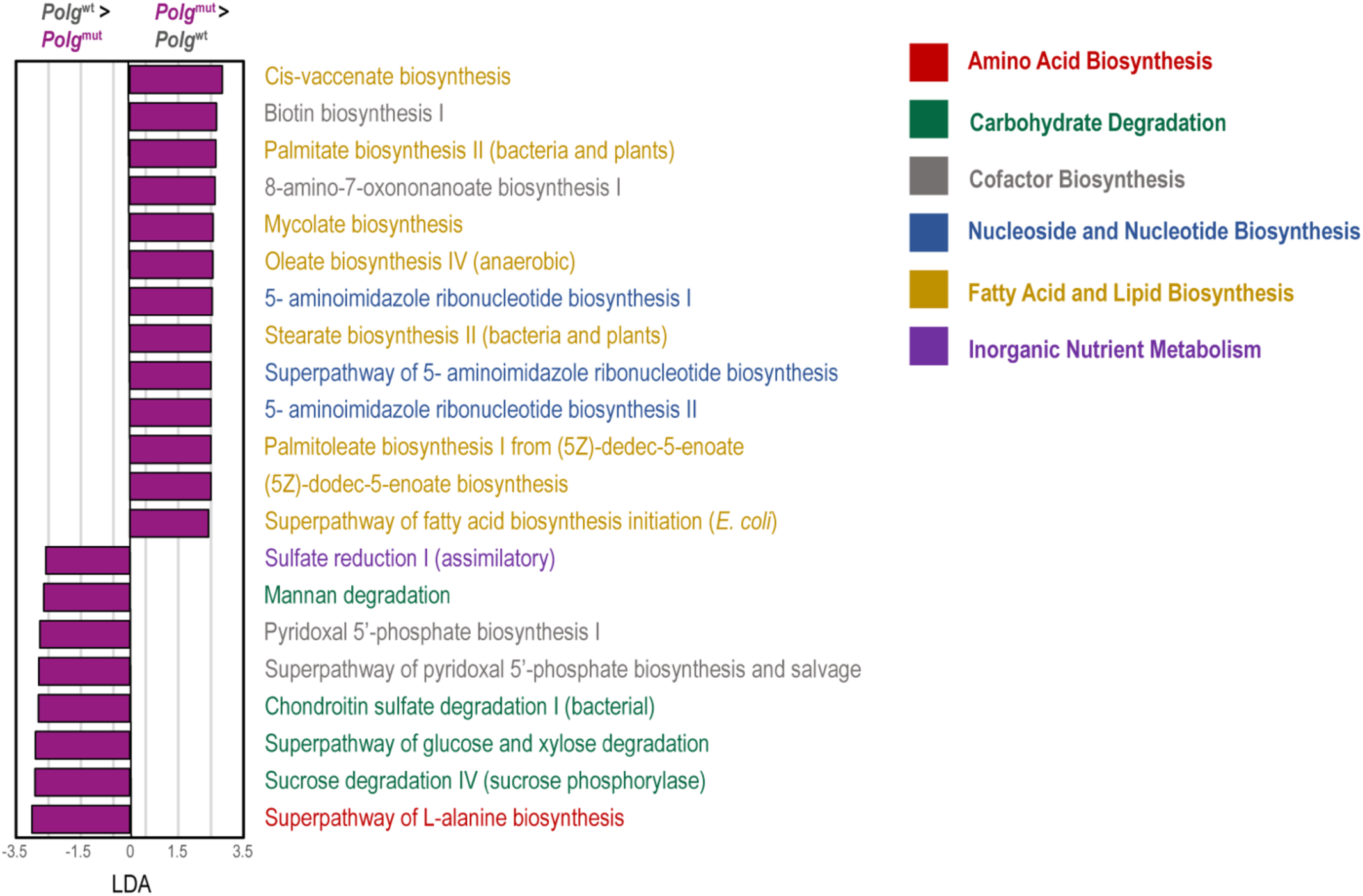
Predictive metabolic profiling of fecal microbiota in mtDNA-mutator mice. PICRUSt2 prediction of metabolic pathways in fecal metagenomic data of control and *Polg*^mut^ mice (FDR, *q* < 0.05; |SNR| > 1; |LDA| > 2).

**Figure S6.**
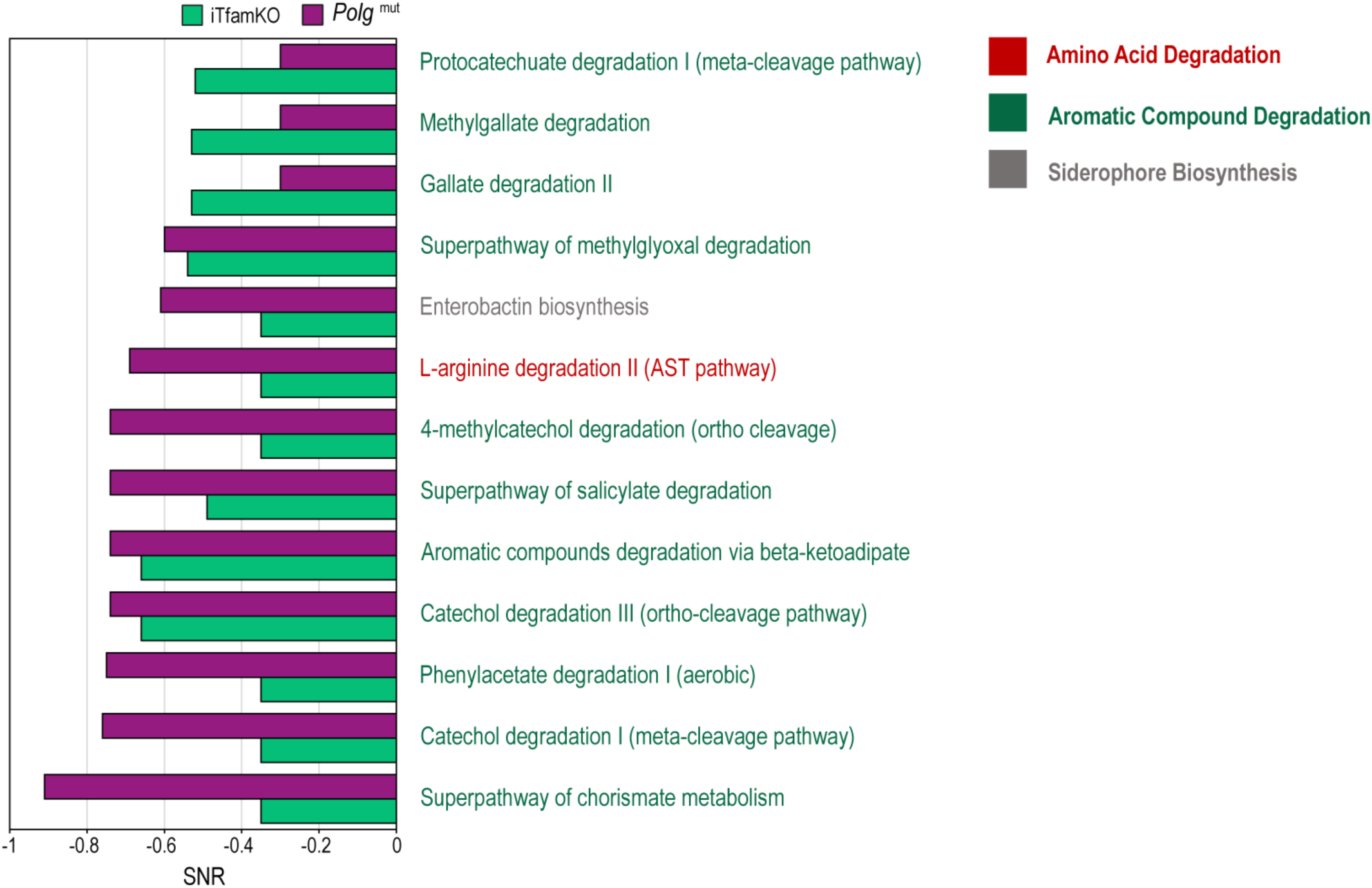
Predictive metabolic profile of intestinal microbiota shared by iTfamKO and mtDNA-mutator mice. PICRUSt2 prediction of shared metabolic pathways in metagenomic data of iTfamKO and *Polg*^mut^ mice (FDR, *q* < 0.05; |SNR| > 0.2; |LDA| > - 1; |fold change| > 0; maximal abundance > 0).

**Figure S7.**
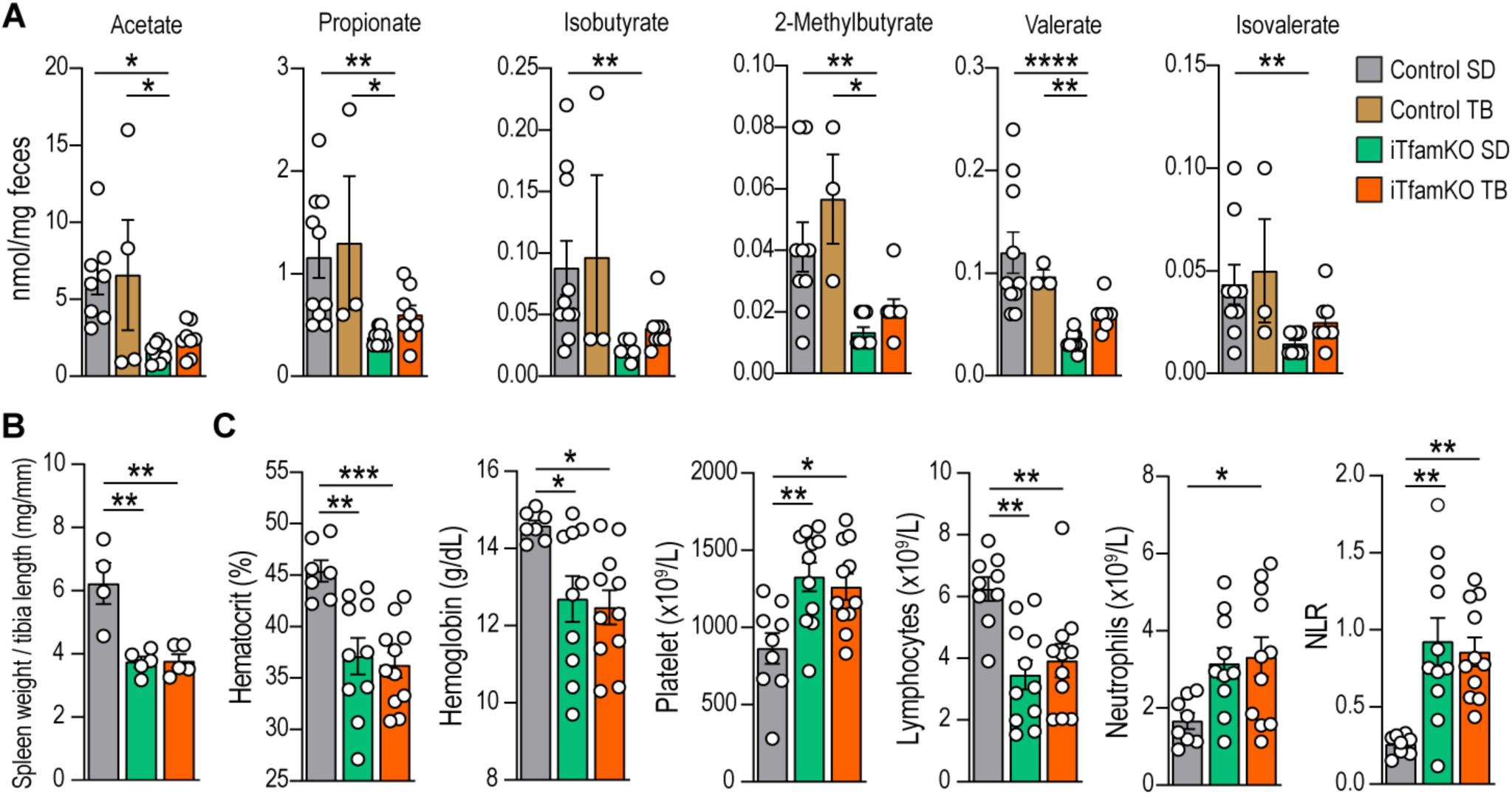
Analysis of SCFAs and hematological parameters in iTfamKO mice after tributyrin administration. (**A**) Quantification of short-chain fatty acids in the feces of control and iTfamKO mice fed either s standard diet (SD) or a 10% tributyrin-supplemented diet (TB) (*n* = 3 to 10). (**B**) Quantification of spleen weight normalized to tibia length (*n* = 4 to 5). (**C**) Quantification of hematological parameters related to red blood cells, platelets, and white blood cells (*n* = 7 to 11). Data are shown as means ± SEM, where each dot is a biological sample. *P* values were determined by one-way analysis of variance (ANOVA) with Tukey’s multiple comparisons test. \**P* ≤ 0.05; \*\**P* ≤ 0.01; \*\*\**P* ≤ 0.001; and \*\*\*\**P* ≤ 0.0001.

**Figure S8.**
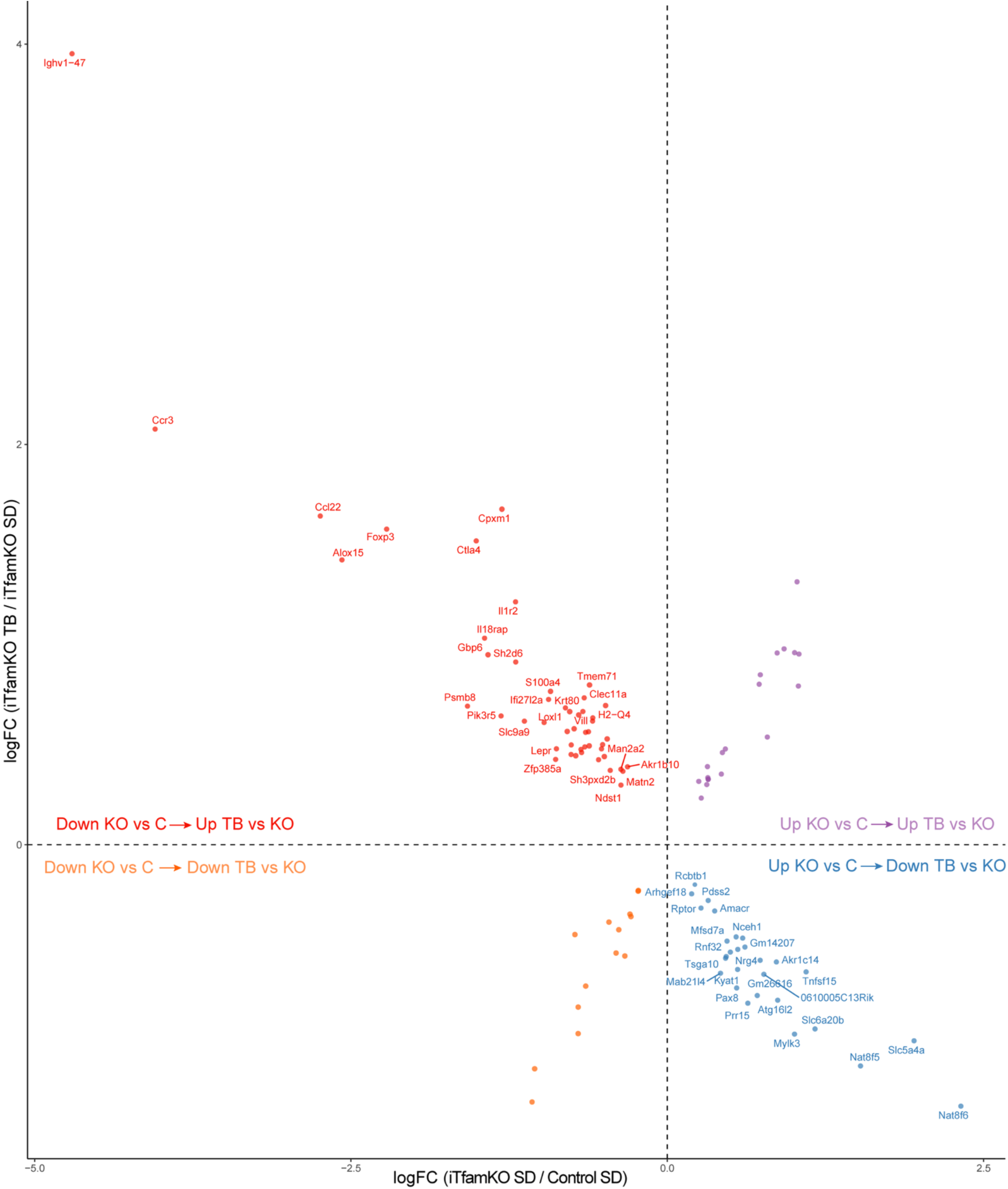
Differentially expressed genes in the small intestine of iTfamKO mice after tributyrin administration. Four-quadrant scatter plot depicting differentially expressed genes in the ileum RNA-sequencing analysis of iTfamKO mice fed either a standard diet (SD) or a 10% tributyrin-supplemented diet (TB).

**Figure S9.**
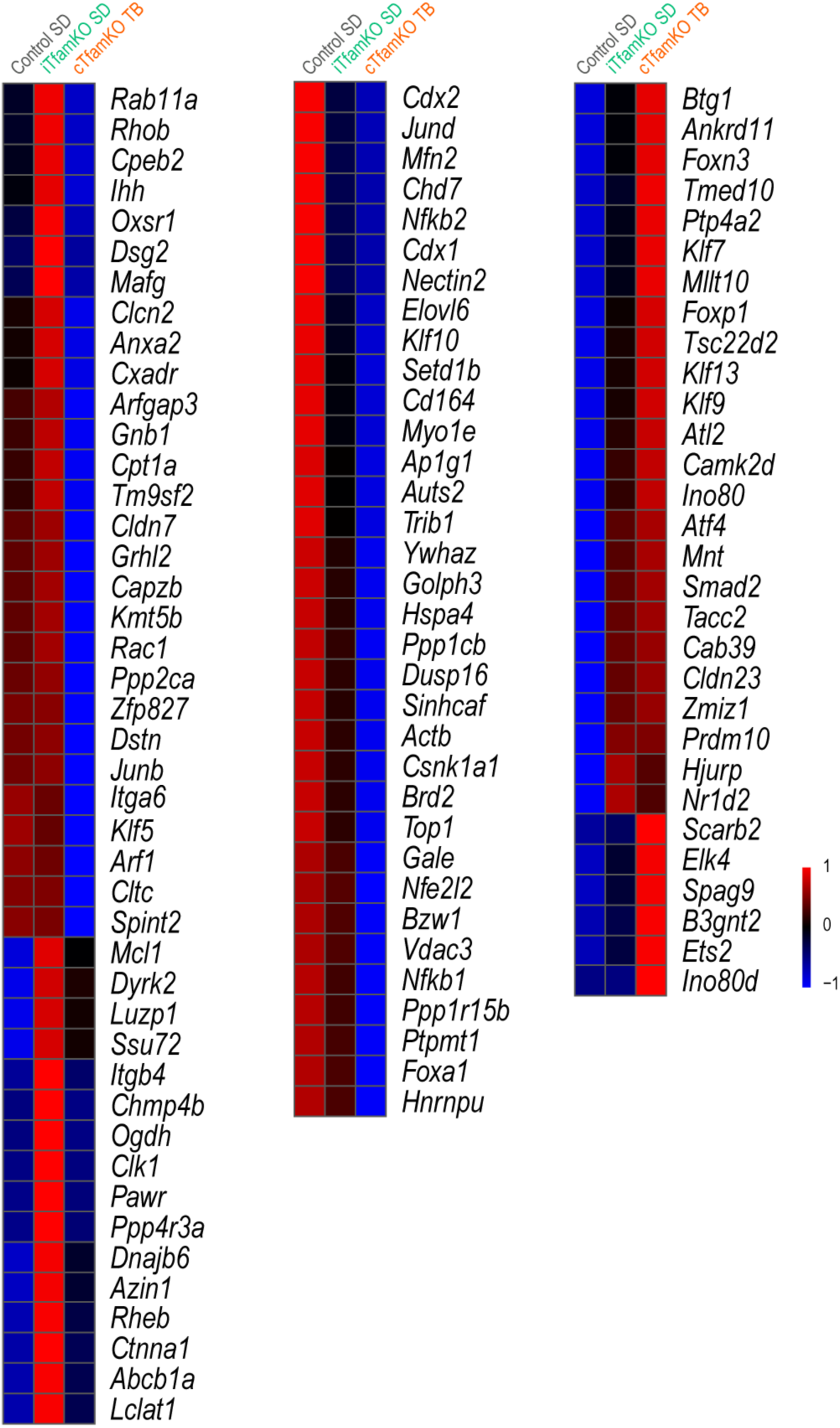
Expression of H3K27bu-dependet genes in the RNA-sequencing analysis of the small intestine in iTfamKO mice after tributyrin administration. Heatmap depicting expression of H3K27bu-dependent genes in the ileum RNA-sequencing analysis of iTfamKO mice fed either a standard diet (SD) or a 10% tributyrin-supplemented diet (TB).

